# Revealing Evolutionary Signals Across Landscapes: A Scalable Genetic Diversity Index for Multi-Species Analysis

**DOI:** 10.1101/2025.06.03.657643

**Authors:** Piyal Karunarathne, Marvin Klümpen, Laura E. Rose

**Author notes:** Marvin Klümpen, Laura E. Rose.

## Abstract

Geographic patterns of community-level genetic diversity provide key information about evolutionary processes and are increasingly relevant for conservation planning under rapid environmental change. Yet, continental-scale, multi-taxon assessments of genetic diversity remain rare, largely due to difficulties in integrating heterogeneous datasets across species and markers. Here, we develop a scalable framework to quantify and map nucleotide diversity (*π*) across co-occurring taxa using georeferenced sequence data. We compiled and harmonized genetic metadata for European vascular plants and calculated site-level *π* for individual species. By spatially aggregating values across taxa, we derived the Genetic Diversity Index (GDI), designed to capture shared geographic structure in intraspecific variation while minimizing species-specific noise. We evaluated robustness using simulations and rarefaction procedures to assess sensitivity to sampling intensity and taxonomic composition. Using ∼630,000 sequences from 1,860 species, GDI identified consistent hotspots of genetic diversity in the Anatolian Peninsula, Southern Iberia, and the Eastern Alps. These regions coincide with areas widely recognized as long-term refugia and biogeographic transition zones. GDI showed weak relationships with species richness and phylogenetic diversity, indicating that it captures a distinct dimension of biodiversity linked to demographic persistence and evolutionary history. In addition, GDI increased significantly with late-Quaternary climate stability, supporting the expectation that regions experiencing reduced climatic fluctuation tend to retain higher levels of genetic variation. Our results demonstrate that publicly available sequence data can be synthesized into reproducible, large-scale estimates of community-level diversity. The GDI provides a practical tool for integrating genetic information into biodiversity assessments and for identifying regions where evolutionary potential may be concentrated. This framework opens new opportunities for incorporating genomic resources into conservation prioritization and macroecological research.

## Introduction

Most biodiversity assessments rely on taxonomic metrics such as species richness or endemism to identify hotspots and set conservation priorities (Miller et al., 2018). However, the processes governing genetic diversity—mutation, gene flow, drift, selection, and demographic history—are largely decoupled from species count or local abundance (Taberlet et al., 2012; Toczydlowski et al., 2021; Hoban et al., 2022; Sills et al., 2020). Consequently, regions rich in species may not coincide with areas of high intraspecific variation, an essential dimension of biodiversity that determines species’ evolutionary potential, adaptive capacity, and long-term persistence under environmental change (Frankham, 2005; Hoffmann et al., 2021).

While species richness maps where biodiversity is concentrated, it offers little insight into the evolutionary and demographic processes that generate and sustain it. In contrast, multi-species patterns of genetic diversity capture these underlying dynamics by integrating intraspecific variation across taxa (Vellend and Losos, 2003; Petit and Hampe, 2006). When aggregated across taxa, intraspecific genetic variation captures legacy signals of population persistence, connectivity, and isolation—features that remain largely invisible to species richness or even phylogenetic diversity metrics, which reflect lineage composition rather than population-level history (Whiting et al., 2024; Bagnoli et al., 2016). Such cross-taxonomic integration of genetic data thus exposes the hidden facets of biodiversity shaped by long-term climatic stability, range dynamics, and local adaptation, offering a powerful lens for identifying landscapes that have repeatedly fostered genetic resilience across lineages (Hoban et al., 2022; Luikart et al., 2019). These insights provide the foundation for our hypothesis that large-scale spatial concordance in intraspecific genetic diversity marks regions of enduring evolutionary stability and adaptive potential.

Existing approaches for large-scale assessments of genetic diversity typically rely on broad ecological or phylogenetic proxies, rather than direct measures of intraspecific variation. These include the use of phylogenetic diversity and related metrics that infer evolutionary history from species relationships (Faith, 1992; Tucker et al.), and environmental or trait-based indices that approximate adaptive capacity under climate change (Hoffmann and Sgrò, 2011). While informative, such proxies do not capture realized patterns of standing genetic variation within species. Over the past decade, advances in high-throughput sequencing, DNA barcoding, and open data policies have vastly expanded public genetic repositories such as GenBank, EMBL-EBI, and DDBJ (Kodama et al., 2012), enabling large-scale analyses of intraspecific diversity beyond traditional single-species studies that required dense, costly sampling (Henle et al., 2014). Recent macrogenetic studies have begun to exploit such data (Paz-Vinas et al., 2018; Blanchet et al., 2020). For example, De Kort et al., 2021 synthesized selected plant and anmal populations worldwide to show that life history, geography, and contemporary climate jointly shape broad-scale patterns of genetic diversity across plants and animals. However, standardized, cross-taxon frameworks for spatially mapping multi-species genetic diversity are still lacking (Troudet et al., 2017). The framework we introduce here addresses these gaps and complements existing approaches by providing a spatially explicit, empirically derived measure of intraspecific nucleotide diversity aggregated across taxa.

The central premise is that regions maintaining high intraspecific genetic diversity across multiple co-occurring species represent evolutionary refugia—areas that have historically buffered populations from environmental change and therefore retain high adaptive potential for the future (e.g., Hewitt, 2000; Hampe and Petit, 2005). Leveraging publicly available sequence data, our approach calculates nucleotide diversity (*π*) for individual accessions and aggregates these values by geographic location to derive a composite measure of site-level diversity, the Genetic Diversity Index (GDI). While *π* itself captures the balance of mutation, drift, and demography within populations, its spatial aggregation across taxa enables the identification of regions that consistently foster evolutionary resilience across lineages, which we interpret as the capacity of populations to maintain standing genetic variation through time. Here, we evaluate whether regions of high GDI are associated with long-term climatic stability, environmental heterogeneity, or historical connectivity—conditions widely proposed to promote the accumulation and maintenance of genetic variation across taxa (Stewart et al., 2009; Alsos et al., 2012). By quantifying these relationships, GDI provides a macro-evolutionary lens for identifying landscapes where adaptive capacity may be concentrated, revealing patterns that are not evident from species richness or phylogenetic diversity alone.

We applied this framework to European vascular plants, integrating data from 623,721 accessions to map large-scale patterns of intraspecific genetic diversity. We then compared these patterns with taxonomic and phylogenetic diversity, revealing four genetic diversity hotspots largely decoupled from traditional biodiversity metrics. Using simulation analyses, we tested the robustness of our approach under varying sampling intensities and diversity scenarios, resulting in three optimized indices of genetic diversity. Together, these results demonstrate both the feasibility of harnessing public genomic data for continental-scale biodiversity assessments and the capacity of the Genetic Diversity Index (GDI) to pinpoint regions of exceptional evolutionary significance. By integrating genetic, spatial, and ecological perspectives, GDI provides a scalable tool for incorporating genetic diversity into conservation planning, monitoring efforts, and biodiversity policy frameworks across spatial scales.

## Materials and Methods

We developed an integrated framework to assess genetic diversity patterns of vascular plants across Europe using publicly available species occurrence, distribution, and sequence data. The complete analysis workflow was divided into two main components: **(i)** data compilation and processing, and **(ii)** genetic diversity index (GDI) calculation and validation. Figure 1 shows the complete workflow, and a detailed description of all computational steps, software, and code used in the analysis is provided in Appendix 1. Below we provide a brief overview of the key methods employed and rationale in each component.

**Figure 1:**
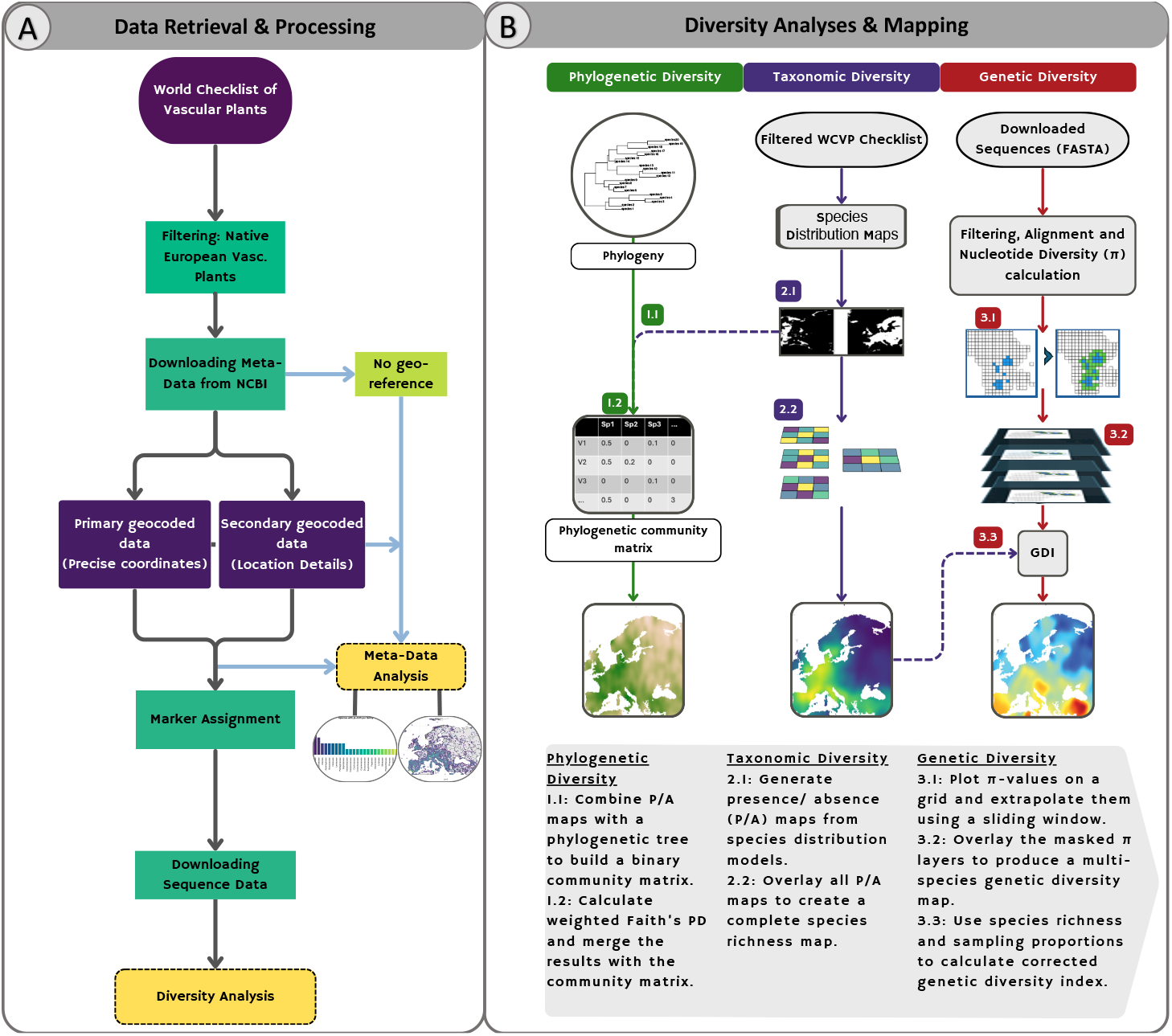
Overview of the data processing and diversity analysis pipeline used to develop the Genetic Diversity Index (GDI). **A. Data retrieval and metadata processing**. We began by compiling a checklist of native and extant European vascular plants using the World Checklist of Vascular Plants (WCVP). Species names were matched against metadata downloaded from the NCBI GenBank database, yielding over 25 million accessions. These were screened for georeferenced records, and a two-tiered classification was performed: primary geocoded sequences contained exact geographic coordinates, while secondary geocoded sequences included place names or locality descriptions. A marker-assignment step was then carried out by searching sequence annotations for ∼50 commonly used plant genetic markers to identify species–marker combinations suitable for alignment and genetic diversity estimation. This metadata-driven filtering process resulted in >600,000 sequences from ∼1,800 taxa with sufficient spatial and genetic representation for downstream analyses. **B. Biodiversity component analyses and spatial mapping**. Three complementary dimensions of biodiversity were assessed at a common 0.25° (∼400 km^2^) spatial resolution: (i) *Taxonomic diversity* was quantified as species richness using stacked presence/absence maps from species distribution maps from Daru (2024). (ii) *Phylogenetic diversity* (PD) was computed as weighted Faith’s PD based on a dated vascular plant phylogeny (Smith and Brown, 2018), with branch lengths integrated via a binary community matrix. (iii) *Genetic diversity* was derived by calculating nucleotide diversity (*π*) from multiple sequence alignments and interpolating these values across space to map multispecies patterns of intraspecific diversity. Corrected GDI values were then obtained by combining *π* with local species richness and sampling proportions. Panels A and B together summarize how public genetic data, taxonomic range maps, and phylogenetic structure were integrated into a unified framework for large-scale genetic diversity assessment across Europe.

### Data Compilation and Processing

A curated list of native and extant plant species from the European region was compiled from authoritative taxonomic sources (e.g., Plants of the World Online database [POWO, 2024]; World Checklist of Vascular Plants [WCVP: Govaerts et al., 2021]) and used to retrieve occurrence records and sequence metadata from the NCBI GenBank database. Representing 9,409 species, we downloaded ∼25 million accessions. A comprehensive metadata analysis on 24,569,450 million records is presented in Appendix 1.

To extract georeferenced genetic accessions relevant for analysis, we first identified and assigned genetic markers by searching sequence descriptions for 55 commonly used plant loci (e.g., *matK, rbcL*, ITS; full list in Appendix 1). This marker-assignment step enabled us to group individual sequences into species–marker combinations suitable for alignment and downstream calculations of genetic diversity. Only species–marker combinations with sufficient representation (at least five accessions) and spatially precise geolocation (i.e., exact coordinates) data were retained, resulting in a high-resolution dataset of 623,721 sequences across 1,851 taxa. These data were used to compute GDI values across a standardized ∼400 km^2^ grid covering Europe. Figure 1A summarizes the data compilation and processing workflow, and a detailed description of all computational steps, software, and code used in the analysis is provided in Appendix 1.

### Genetic Diversity Index (GDI) Calculation and Validation

The core aim of this study was to develop a novel, spatially explicit Genetic Diversity Index (GDI) that quantifies intraspecific genetic variation across communities and geographic space. We used the mean nucleotide diversity (*π*) of all individuals of all species occurring within a given spatial unit (0.25° raster cell: ∼400 *km*^2^) as neutral genetic diversity at a site, which we call “*m*GDI”.

#### Contextual Justification

Averaging nucleotide diversity across co-occurring species is biologically justified under our central assumption that large-scale spatial concordance in intraspecific diversity reflects shared historical and environmental drivers of population persistence. Although individual species may exhibit idiosyncratic peaks of *π* due to demographic or ecological peculiarities, broad-scale patterns of high or low diversity tend to emerge in regions that have acted as long-term refugia, dispersal corridors, or zones of environmental stability influencing multiple taxa simultaneously. In this context, the mean *π* across species does not aim to capture species-specific demographic histories, but rather to extract the common spatial signal of evolutionary resilience that arises when many lineages have independently retained elevated genetic variation in the same landscape units. Consequently, localized discordance among species (e.g., species A peaking at Site 1, species B at Site 2) contributes to noise but not systematic bias, while spatially concordant trends reinforce the regional signature of long-term evolutionary stability. The resulting mGDI therefore provides a robust, community-level summary of where evolutionary potential is concentrated across the landscape—an interpretation consistent with the hypothesis that high GDI regions correspond to long-term climatic stability, environmental heterogeneity, or historical connectivity among populations.

To do this, we first calculated the per-individual average nucleotide diversity (*π*) for each species, based on all pairwise comparisons among its sequences within each species–marker combination (e.g., *Abies alba*–rbcL). For species represented by multiple markers, we retained only the species–marker combination with the widest spatial representation, alignining only the homologous marker sequences to ensure consistent and geographically representative estimates of diversity. Multiple sequence alignments were performed using MAFFT (Katoh and Standley, 2013), with settings optimized for maximum accuracy (i.e., G-INS-i). All alignments were manually inspected to ensure proper alignment quality and to correct any misalignments. We applied a threshold of 80% sequence coverage to filter out poorly aligned or incomplete sequences.

The individual nucleotide diversity was calculated as

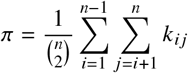

where *n* is the number of sequences and *k*_*ij*_ is the proportion of nucleotide sites at which the *i*^th^ and *j*^th^ sequences differ (Nei and Li, 1979). The resulting per-individual *π* values were then assigned to spatial grid cells according to the geographic coordinates of each accession, yielding a three-dimensional matrix of *π* values across all species (i.e., *x* = longitude, *y* = latitude, *z* = species). Finally, we computed the mean nucleotide diversity (*m*GDI) for each grid cell as

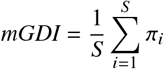

where *S* is the number of species with genetic data in that grid cell and *π*_*i*_ is the per-individual mean nucleotide diversity of species *i*.

To account for potential biases arising from uneven species sampling and varying species richness across grid cells, we developed three complementary but mutually exclusive variants of the GDI:

i. weighted genetic diversity index (*w*GDI),–the mean nucleotide diversity weighted by species richness and sampling proportion, calculated as

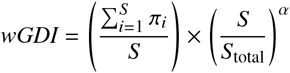

where *S* is the number of species sampled at the site, *S*_total_ is the total number of species predicted to occur at that site, and α is a scaling factor to adjust for sampling bias and can be tuned based on empirical data (default α = 1),
ii. corrected genetic diversity index (*c*GDI)–a regression-based correction of mean nucleotide diversity to account for uneven sampling and species richness effects; First we fit a linear model to predict GDI as a function of sampling proportion at each site, and then the corrected GDI is calculated using the model residuals; cGDI removes the linear dependency between observed GDI and sampling proportion.

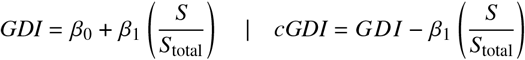
iii. cumulative genetic diversity index (*s*GDI)–the sum of nucleotide diversity across all species at a site, calculated as

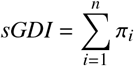

where *π*_*i*_ is the nucleotide diversity of species *i* at that site.

We also performed a rarefaction analysis to standardize GDI values across grid cells. This involved subsampling sequences within each grid cell to a common number of accessions (e.g.20% of total sampling) and recalculating GDI for each subsample. This approach allowed us to assess the robustness of GDI estimates to sampling intensity and ensured comparability across sites with differing data availability. These GDIs represent a flexible and scalable approach to quantifying spatial patterns of genetic variation in different ecological and sampling contexts. A detailed description of the GDI calculation, including the underlying equations, further extensions, and computational steps, is provided in Appendix 1. Special use scenarios for each of these indices are described in the Results and Discussion sections.

To contextualize spatial patterns of GDI, we derived comparable estimates of taxonomic and phylogenetic diversity following established workflows (Daru et al., 2020; Daru, 2024). Species richness was calculated as the number of vascular plant species occurring in each 0.25° (∼400km^2^) grid cell using distribution maps for 12,571 native taxa from Daru (2024), which integrate expert-curated occurrences, species distribution models, and machine-learning extrapolations to provide continent-wide coverage. Phylogenetic diversity was quantified using Faith’s PD (Faith, 1992) based on the dated vascular plant phylogeny of Smith and Brown (2018). The tree was pruned to match the species set represented in the richness layer, and values were standardized to account for variation in species numbers among cells. Both layers were generated at the same spatial grain as GDI to enable direct comparison among taxonomic, phylogenetic, and genetic dimensions of biodiversity (Fig. 3B–C).

In our analysis, these served as comparative layers to evaluate how genetic diversity aligns with broader dimensions of biodiversity. The GDI’s performance and robustness were validated through permutation-based resampling, correlation assessments, and large-scale simulations, confirming that the corrected indices effectively mitigate sampling biases and species richness effects. The resulting maps highlight spatial heterogeneity in genetic diversity and provide a basis for integrative conservation planning across ecological and evolutionary scales. A detailed explanation of all computational steps and validation procedures is provided in Appendix 1.

### Association between climate stability and GDI

To test whether regions of high multispecies genetic diversity coincide with long-term climatic persistence, we quantified late-Quaternary climate stability and evaluated its relationship with the Genetic Diversity Index (GDI). We used bioclimatic variables from the WorldClim CCSM4 dataset, which provides high-resolution reconstructions of temperature and precipitation across four key time slices: the Last Interglacial (LIG∼140 kyr ago), Last Glacial Maximum (LGM∼12 kyr ago), Mid-Holocene (∼4 kyr ago), and present. These time slices capture major climatic transitions that have shaped species distributions and genetic diversity patterns in Europe. For each grid cell, temperature and precipitation stability were calculated as the negative standard deviation of bioclimatic variables across the four time slices. Higher values therefore represent more stable climatic conditions through time. Stability layers were resampled to the GDI grid and standardized (z-scores). We first assessed bivariate associations using Pearson and Spearman correlations. We then fitted linear models with standardized predictors:

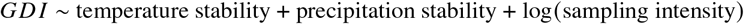

to evaluate effect sizes while accounting for variation in data availability. Because spatial autocorrelation can inflate significance, we further fitted generalized additive models (GAMs) including a two-dimensional smooth of geographic coordinates.

## Results

### Insights from the metadata-analysis of European Vascular Plant Sequences

Our analysis revealed that of Europe’s 26,242 native vascular plant species, 9,409 (∼38%) are represented by ∼25 million GenBank accessions, of which ∼13% are georeferenced. Primary coordinates were available for 280,979 accessions (∼1.2%) and secondary locality data for 2.77 million (∼11.7%; Fig. 1). Among sequenced species, 2,827 (∼30%) had primary and 7,143 (~72%) had secondary geolocation data, together representing 7,512 species (∼77% of sequenced taxa; see Appendix Fig. S6 for detailed analysis). After quality filtering—retaining only species– marker combinations with ≥5 accessions and the best-represented marker per species—the dataset comprised 1,860 species across 629,175 accessions (mean ≈94 sequences per species–marker).

Taxonomic representation was uneven: large families such as Asteraceae (12%), Rosaceae (14%), and Ranunculaceae (12%) were underrepresented, whereas smaller families (7–1,094 species) averaged ∼45% coverage (see Appendix Fig. S7). Spatially, georeferenced sequences were highly clustered, with 1–2,138 sequences per ∼20 km grid cell (mean = 135), and hotspots around major cities such as Antwerp, Prague, Ankara, and Madrid (Appendix Fig. S10). Detailed taxonomic and spatial distributions are provided in Appendix Figs. S6–11.

### Genetic Diversity Patterns in European Vascular Plants

Nucleotide diversity (*π*) across all the analyzed species exhibited substantial variation, ranging from 0 to 0.2415. The minimum *π* range per species spanned 0 to 0.0971, while the maximum ranged from 0.0143 to 0.2286, highlighting notable heterogeneity in genetic variation among European vascular plants.

Across species, geographic range size was positively associated with the variability of nucleotide diversity. Species occupying more grid cells exhibited broader ranges of *π* values (Spearman ρ = 0.31, p < 0.001), indicating that widespread taxa tend to encompass both high- and low-diversity populations. Similarly, species occurring across a greater number of ecoregions showed higher mean *π*, suggesting a link between environmental heterogeneity and within-species genetic variation. Individual examples such as *Sisymbrium officinale* (*π*: 0.07–0.23) and *Pancratium maritimum* (*π*: 0–0.245×10^−4^) illustrate these broader trends. Interestingly, several species with high *π* values also exhibited disjunct distributions, including *Haplophyllum buxbaumii, Cerastium grandiflorum*, and *Veronica scardica*, possibly reflecting long-term isolation or historical refugia.

At the family level, *Asteraceae* and *Poaceae* included the highest number of species with elevated nucleotide diversity (*π* > 0.05), reflecting many species with substantial genetic variation. However, due to their large overall species richness, families such as *Poaceae* and *Fabaceae* also contained a considerable number of species with low genetic diversity (*π* < 0.05). This pattern highlights that, while large families harbor many genetically diverse species, they simultaneously include numerous species with limited diversity, resulting in substantial heterogeneity in genetic variation even within major plant families (Fig. 2). Higher taxonomic levels (orders and clades) also exhibited significant variation in nucleotide diversity (Fig. 2A). For example, orders such as *Fagales* and *Solanales* showed higher mean *π* values compared to orders like *Poales* and *Lamiales*, indicating that evolutionary history and life-history traits at these taxonomic levels may influence patterns of genetic diversity.

**Figure 2:**
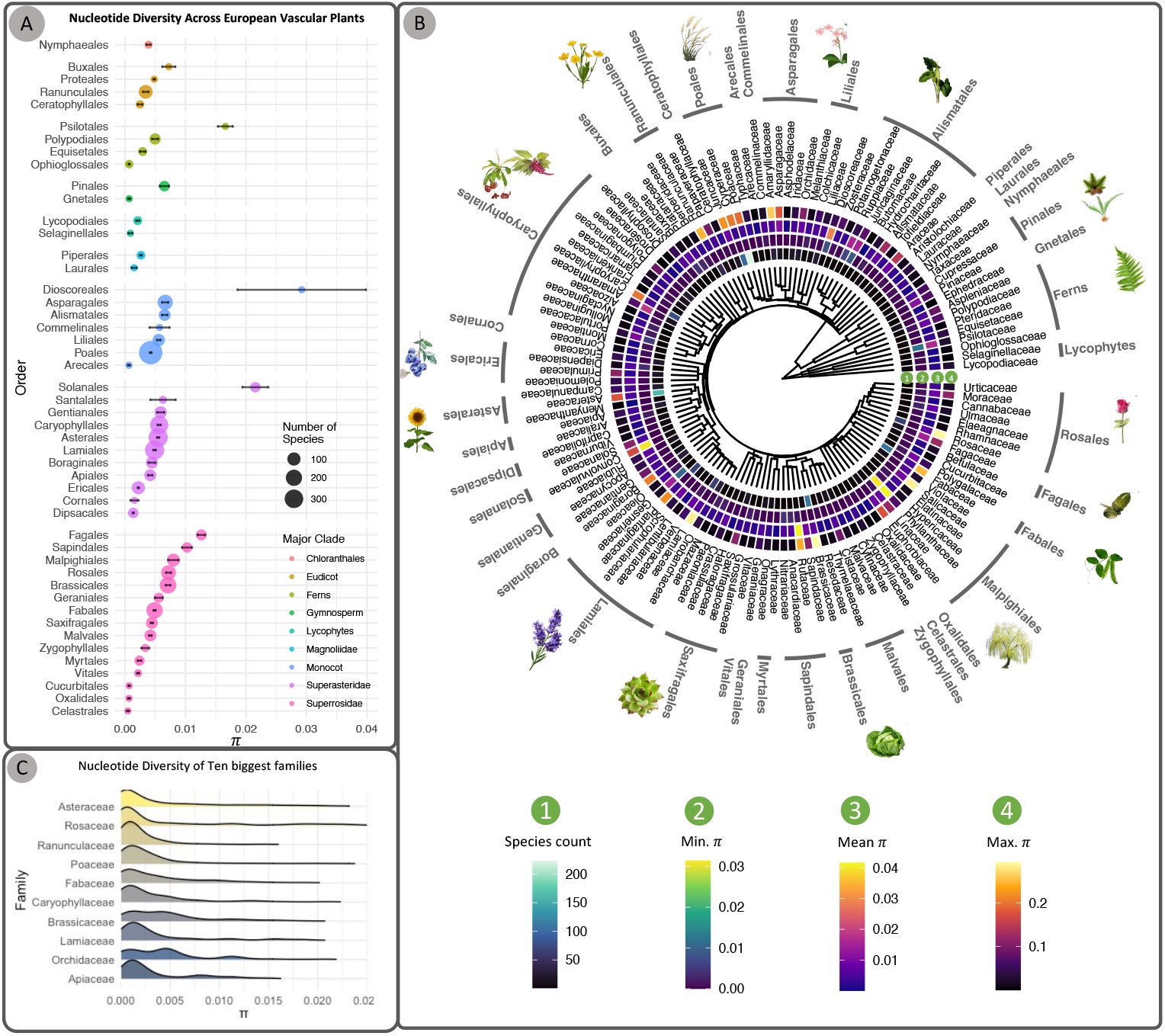
Nucleotide diversity variation among higher taxonomic levels of vacular plants in Europe. **A.** Pyramid plot showing the distribution of nucleotide diversity (*π*) values across the major orders of European vascular plants. Error bars represent standard errors, extending from the order mean. Circle sizes correspond to the number of species sampled within each order. Orders are arranged from lowest to highest mean *π* values. **B**. Family-level phylogenetic tree of all vascular plants in Europe, constructed using the R package V.Phylomaker2 (Jin and Qian, 2019), which uses an extended version of GBOTB tree from Smith and Brown (2018). The heatmap surrounding the tree visualizes (1) number of species sampled, (2) minimum *π*, (3) mean (*π*), and (4) maximum *π* for each family, with colors gradients. **C**. Kernel density plots illustrating the distribution of nucleotide diversity (*π*) values for species within the largest plant families in Europe.

### Spatial Patterns of Genetic Diversity in European Vascular Plants

Our genetic diversity index (GDI) revealed a highly heterogeneous spatial distribution of genetic diversity across Europe (Fig. 3 and Appendix Fig. 10). Based on these patterns, we identified four major hotspots and several low-diversity zones, each reflecting known biogeographic and ecological gradients. Table 1 summarizes these patterns, showing the GDI range, mean values, and number of species sampled for each major region. These values are the corrected GDI reflecting combined nucleotide diversity across species. Unless otherwise noted, all mentions of GDI hereafter refer to the corrected version.

**Figure 3:**
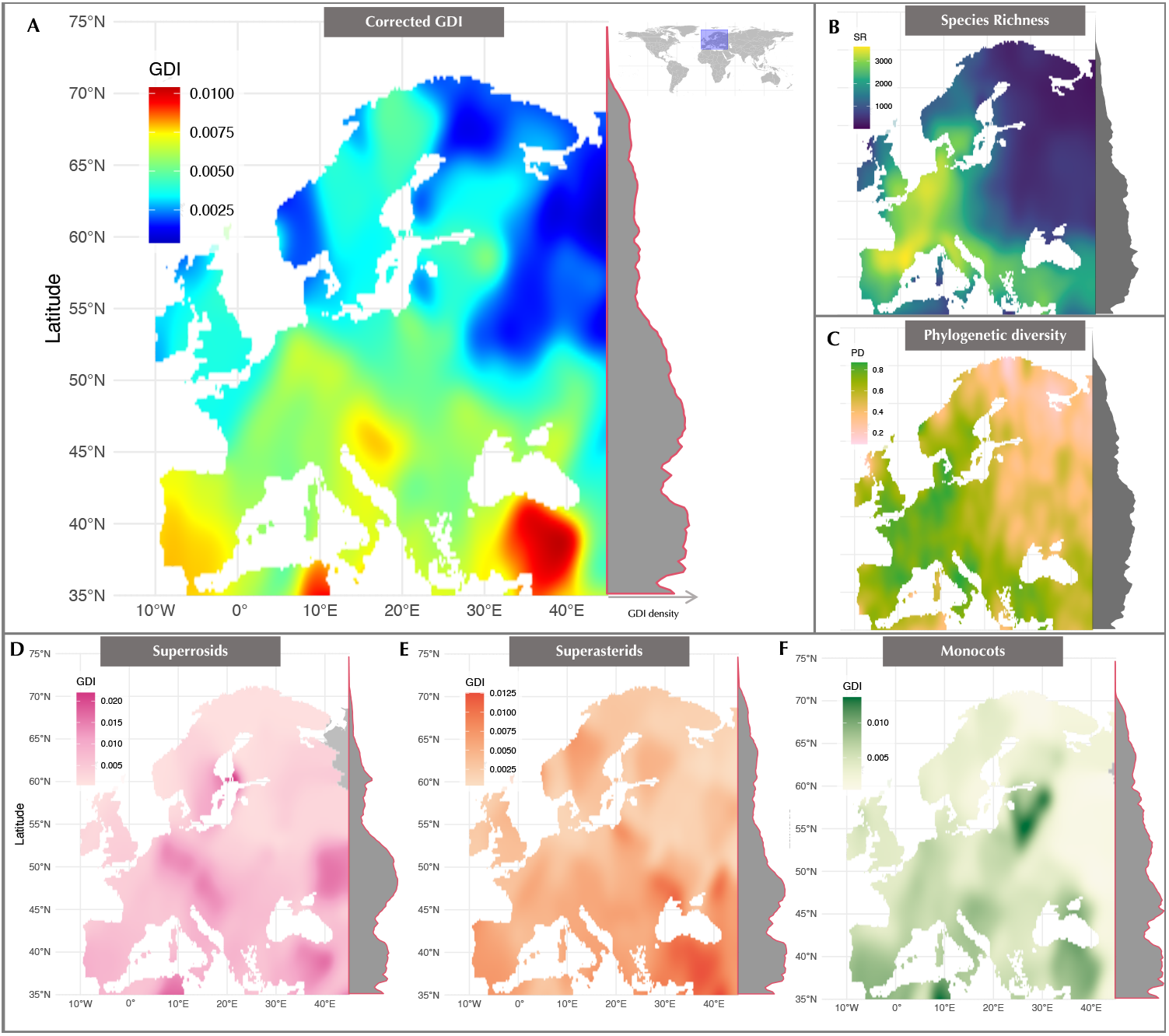
Spatial distribution of genetic (A, D, E, F), taxonomic (B), and phylogenetic (C) diversity of vascular plants across Europe. Genetic diversity is represented by the Genetic Diversity Index (GDI) developed in this study: corrected GDI (*c*GDI; mean nucleotide diversity, *π*, across all species at a site corrected for sampling bias). Nucleotide diversity *π* measures the average number of nucleotide differences per site between two sequences within a species. Taxonomic diversity (B) is species richness (total number of species), calculated from species distribution models, curated datasets, and machine-learning extrapolations from Daru (2024). Phylogenetic diversity (C) is Faith’s D standardized for species richness (Chao et al., 2010). Genetic diversity distribution of the three biggest vascular plant clades of Europe, Superrosids (D), Superasterids (E), and Monocots (F) shows group-specific contribution to complete GDI. Grey polygons along the right side of each map illustrate the latitudinal density distribution of diversity values. Regional delineations follow TDWG ecological categories (Brummitt et al., 2001) and do not correspond to political boundaries.

**Table 1:**
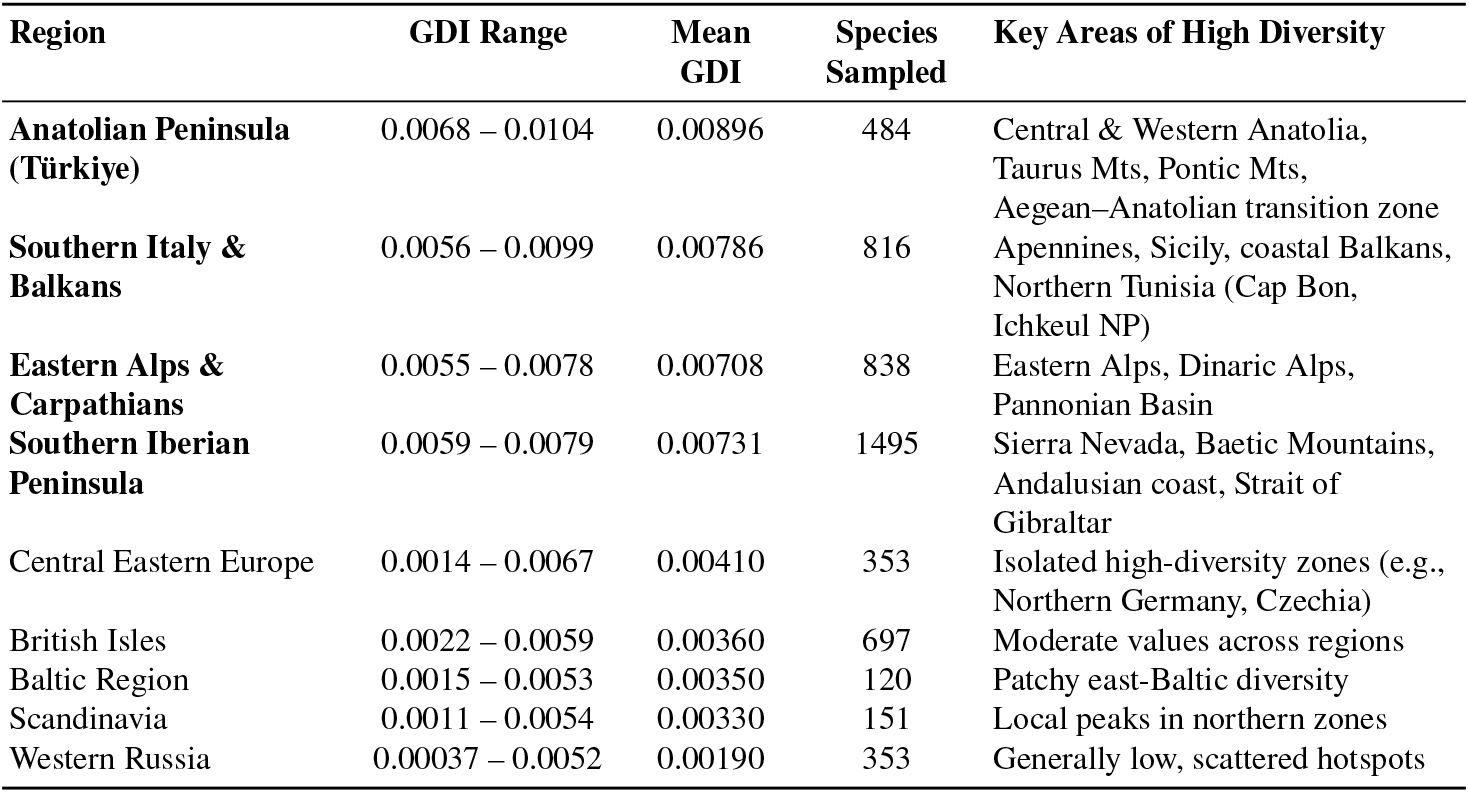
Summary of Genetic Diversity Index (GDI) and Sampling Metrics Across European Regions. Genetic diversity hotspots are in **bold** font. Sampled species are the number of species with usable georeferenced data in each region. The mean GDI is the average GDI value across all sampled species in each region. Key areas of high diversity are the regions within each area that exhibited the highest GDI values.

Among the hotspots, the **i) Anatolian Peninsula** stood out with the highest average and maximum GDI values. This region includes topographically complex mountain systems and historical refugia, supporting high endemism and long-term population persistence. Similarly, elevated diversity in **ii) Southern Italy, the Balkans**, and parts of North Africa reflects stable climatic conditions and complex biogeographic histories. **iii) Eastern Alps and Carpathians** have served as refugia for plants over many glaciation cycles and have high species richness. **iv) Southern Iberian Peninsula** is a contact zone and an ecotone (transition zone) connecting Mediterranean, Atlantic and North African flora with its heterogeneity of topography, microclimate and endemism. Notably, the GDI patterns of the three largest vascular plant clades, Superrosids, Superasterids, and Monocots, exhibit subtle but distinct differences from the overall species-level GDI, suggesting that individual clades contribute differentially to the spatial structuring of genetic diversity across Europe (see Fig. 3D-F).

In contrast, Scandinavia, Western Russia, and Baltic Europe showed low genetic diversity, likely shaped by postglacial recolonization and reduced habitat complexity. However, localized peaks in northern and eastern areas suggest microrefugia or contact zones among divergent lineages.

### Genetic Diversity Index Enables Robust and Independent Comparisons

To evaluate the robustness and distinctiveness of our Genetic Diversity Index (GDI) in capturing spatial genetic diversity patterns, we compared it with two widely used biodiversity measures: taxonomic diversity (species richness) and phylogenetic diversity (PD). Although the genetic dataset comprises only the subset of species with available sequence data, we used complete species richness and PD maps to represent the full ecological and evolutionary diversity of each site. This approach is justified because our simulation analyses demonstrated that GDI estimates remain highly accurate when genetic data cover at least 25% of the local flora, with relative errors below 10% and predicted GDI values exceeding 95% of those derived from complete sampling (details in section below). Ninty eight percent of our sites (raster cells) had at least 25% species representation, and, accordingly, we calculated a site-level confidence metric (*C*) that integrates local species richness and sampling proportion to quantify the reliability of each GDI estimate. Thus, while the underlying datasets differ in species composition, the comparisons presented here are robust, interpretable, and standardized by an explicit confidence framework.

### Taxonomic and Phylogenetic Diversity Patterns

Taxonomic diversity showed a gradual decline from south to north, with the Mediterranean region harboring the highest species richness, averaging around 3,200 species per grid cell. Notably, the Pyrenees Mountains, Alps, and Apennine Mountains emerged as taxonomic hotspots, with mean species richness values of 3,424, 2,942, and 2,730, respectively (Fig. 3). Additionally, temperate biomes such as the western European broadleaf forests and Atlantic mixed forests exhibited substantial richness (∼3,120 species), consistent with previous estimates based on large scale plot analyses (e.g., Sabatini et al., 2022) and machine learning approaches (e.g., Daru, 2024).

Phylogenetic diversity largely mirrored taxonomic diversity, exhibiting a similar latitudinal gradient (Fig. 3). The two metrics were strongly correlated (Pearson’s r = 0.913; also see Fig. 4F & Appendix. Fig. 8 for non-linear correlation), with high PD values observed in the Pyrenees, Alps, Dinaric Alps, southern Italy, and western Anatolia (average PD = 0.738). Surprisingly, northern Germany and Denmark corresponding to the Baltic and Atlantic mixed forests also exhibited elevated PD (average PD = 0.743), alongside western France and southern British Isles (PD = 0.702), highlighting the contribution of relict or phylogenetically distinct species in these regions.

**Figure 4:**
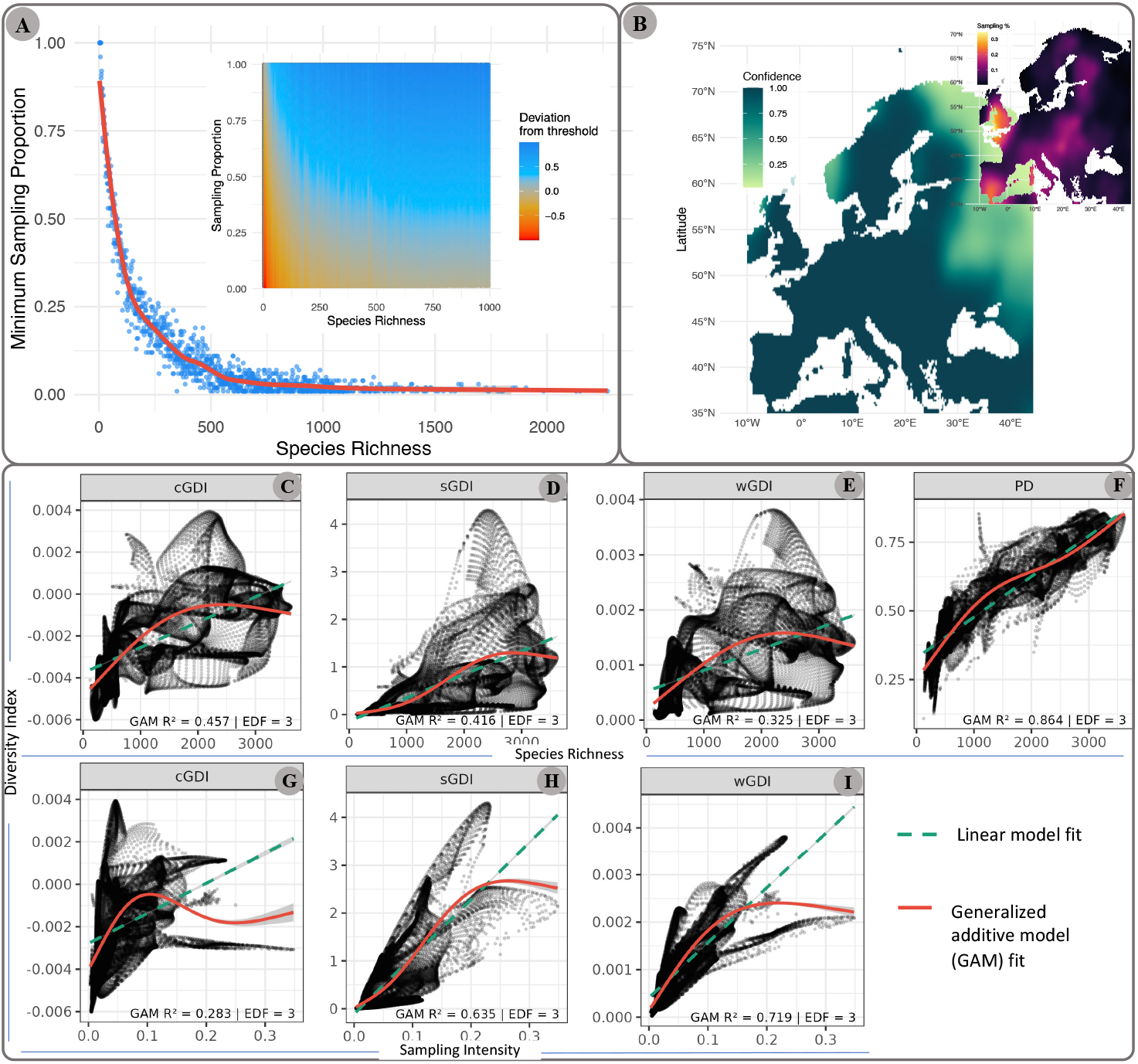
Simulation-based calibration and empirical validation of GDI robustness. (**A**) Simulation-based relationship between species richness (*R*), sampling intensity (*s*), and the relative error of GDI estimates per site. The curve indicates the minimum sampling threshold (*s*^∗^ = *f* (*R*))—the proportion of species that must be sampled at a site for 95% of simulated GDI estimates to deviate by less than 10% from true values. The inset shows the deviation of simulated GDI estimates from their true values across combinations of sampling intensity and species richness, expressed relative to the reliability threshold *s*^∗^. Positive values indicate sampling levels exceeding the threshold (high confidence–blue), negative values indicate insufficient sampling (low confidence–red), and zero marks the minimum sampling proportion required for reliable estimation. (**B**) The main figure shows the site-level confidence *C* in GDI estimates for the empirical dataset, calculated using the simulation-derived function *s*^∗^ = *f* (*R*) and the confidence model described in the Methods. For each site, confidence represents the expected reliability of its GDI estimate given the observed sampling intensity *s* and species richness *R*. Sites with *s* ≥ *s*^∗^ achieve full reliability (*C* = *P*_*R*_), while those with lower sampling are penalized linearly according to their deviation from the threshold. Higher values indicate sampling conditions exceeding the minimum requirement for reliable estimation. The color gradient represents the confidence level (dark green = high confidence, light yellow = low confidence). The inset shows the distribution of sampling intensities across all sites. **C–I**. Linear and non-linear relationships between diversity metrics and species richness (C–F) and sampling intensity (G–I), shown using Linear model (dashed-line) and Generalized Additive Model (GAM) fits with 95 % confidence ribbons (solid-line). Each panel represents a diversity metric: corrected GDI (*c*GDI), cumulative GDI (*s*GDI), weighted GDI (*w*GDI), and Phylogenetic Diversity (*PD*), respectively. Points represents sites. Annotated statistics in each panel indicate the *deviance explained* (proportion of variation accounted for by the model), *GAM R*^2^ (goodness of fit for Gaussian models), and *effective degrees of freedom (edf)*, which describe the degree of non-linearity (edf = 1 ≈ linear; higher values indicate greater curvature).

Despite their high correlation, we retained both metrics because species richness reflects compositional (taxonomic) diversity, whereas phylogenetic diversity incorporates evolutionary relatedness and lineage distinctiveness, providing complementary ecological and evolutionary baselines against which to assess the performance and independence of GDI.

### Robustness to sampling intensity and species-specific baselines

We recognize that nucleotide diversity varies among species because of differences in mutation rate, demography, and life history. Moreover, grid cells differ in the number of available sequences. To evaluate whether the spatial pattern of GDI could be driven by unequal representation of taxa or by species with intrinsically high diversity, we implemented a rarefaction-based normalization in which 20% of records per cell were repeatedly resampled 10,000 times prior to aggregation. GDI values derived from rarefied datasets were highly concordant with those obtained from the complete dataset (Pearson r = 0.93; Spearman *ρ* = 0.91; p ≪ 0.001), indicating that the spatial distribution of diversity is robust to unequal sampling intensity. Appendix Fig. S2 shows the rarefied GDI map. These results indicate that the observed geographic structure is not an artefact of uneven sampling effort or dominance by particular taxa, but instead reflects a shared spatial signal across multiple lineages.

### GDI Captures a Distinct Dimension of Biodiversity

Phylogenetic diversity and species richness were strongly correlated (Pearson’s *r* = 0.913). In contrast, GDI exhibited near-zero correlations with both richness (*r* = 0.0013 for mGDI; *r* = −0.0011 for cGDI) and sampling intensity (*r* = 0.00023 and *r* = −0.0001, respectively) (Fig. 4C–I; Appendix Figs. S3–S4).

To evaluate these relationships, we fitted both linear models and generalized additive models (GAMs). The GAM framework allowed flexible, non-linear responses while accounting for spatial structure through smooth functions. GAMs revealed a moderate association between richness and GDI, with species richness explaining up to 46% of the deviance in some formulations (Fig. 4). This indicates that while GDI is not linearly related to richness, broad-scale spatial gradients in the two metrics are partially aligned. Importantly, a substantial proportion of variation remains unexplained, demonstrating that GDI captures additional information beyond species counts or phylogenetic breadth.

In contrast to traditional diversity measures, which are often influenced by sampling intensity or evolutionary relatedness, the GDI reflects the intraspecific genetic variation accumulated through demographic history, mutation, gene flow, and past environmental changes. These discrepancies are biologically meaningful, as GDI captures variation shaped by evolutionary and demographic processes. Regions with high GDI but moderate species richness often correspond to areas of long-term climatic stability or glacial refugia that have maintained deep genetic structure and adaptive potential across taxa. This makes GDI a valuable and complementary tool for biodiversity monitoring, especially in conservation contexts where genetic resilience is a priority.

Our cumulative GDI (*s*GDI) and weighted GDI (*w*GDI) showed modest correlations with sampling intensity but were less influenced by species richness. These indices are most suitable when sampling is relatively even across sites (see Fig. 4G-I). While they offer complementary insights into spatial patterns of genetic diversity, they should be interpreted alongside a broader understanding of genetic variation across taxa and controled sampling conditions (see Discussion for ideal use).

### Assessing Sampling Effects and GDI Robustness

To quantify how sampling intensity (i.e., species sampled/taxonomic diversity) and species richness affect GDI estimates, we conducted simulations across a range of conditions (species richness: 10–10,000; sampling intensity: 1–100%) at 100,000 sites. For each simulation, GDI values were generated based on the empirical distribution of nucleotide diversity across species. Deviations were then calculated by comparing GDI values at partial sampling to the corresponding GDI values at complete sampling (100% of species sampled).

Our results revealed that, for sites with ≥80 species, GDI estimates remained stable above a 25% sampling threshold (Wilcoxon signed-rank test, *p* = 1); for sites with 15–80 species, sensitivity to sampling increased below ∼50% (typically *p* > 0.05); and for species-poor sites (<15 species), significant deviations persisted even at 85% sampling. Based on these results, we assigned a confidence score to each grid cell, depending on species richness and sampling intensity (Fig. 4B).

We define the minimum sampling intensity threshold *s*^∗^ = *f* (*R*) as the proportion required at species richness *R* for 95% of GDI estimates to have a relative error below 10%; this threshold is based on simulations. For observed sampling intensity *s* ≥ *s*^∗^, the confidence in estimation is *P*_*R*_, the empirical reliability at richness *R*. For lower sampling, confidence is penalized linearly by the proportional deviation from the threshold: *s*^∗^ is the minimum sampling proportion required for reliable estimation (e.g., relative error < 10% in 95% of simulations), and *f* (*R*) is a function empirically derived based on simulation.

To describe the expected confidence *C* in a given sampling intensity *s*, we define:

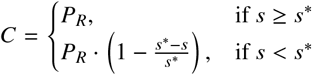

Where: *P*_*R*_: proportion of simulations with error < 10% at richness *R* (i.e., empirical confidence at threshold from the model), and *C*: predicted confidence of accurate GDI estimation at sampling intensity *s* (see Fig. 4B).

Notably, our regression-based correction consistently reduced bias in GDI estimates, outperforming raw simulations and confirming that the corrected GDI is robust to uneven sampling.

### Significance of Genetic Diversity Hotspots and Gaps

The spatial patterns revealed by the Genetic Diversity Index (GDI) highlight unique hotspots of genetic diversity in European vascular plants that are not fully captured by taxonomic or phylogenetic metrics (Fig. 3). While species richness and phylogenetic diversity primarily reflect species turnover and evolutionary distinctiveness, the GDI integrates intraspecific variation across co-occurring species, representing an **emergent property** of biodiversity that arises from the combined effects of microevolutionary processes—mutation, gene flow, drift, and demographic history—acting at multiple hierarchical levels. This emergent nature reflects how aggregated genetic variation across taxa and space produces patterns not predictable from species-level or phylogenetic structure alone. Genetic diversity hotspots frequently coincide with ecological transition zones, topographically complex regions, and known glacial refugia, where long-term population persistence and isolation have fostered high genetic variability. These areas act as reservoirs of adaptive potential, making them critical for long-term resilience under environmental change.

A striking example of this distinction is seen in the Pyrenees Mountains, which exhibit high taxonomic and phylogenetic diversity, indicating strong species turnover and evolutionary divergence. However, the genetic diversity signal (GDI) shifts southward into the Iberian Peninsula, especially in the Baetic and Sierra Nevada regions, where prolonged demographic stability likely contributed to the accumulation of intraspecific diversity across many lineages. Similarly, while the Alpine region displays high species and phylogenetic diversity throughout, the GDI highlights greater diversity toward the Eastern and the Dinaric Alps, regions characterized by sharp ecological gradients and biogeographic transitions. This suggests that intraspecific genetic variation may be highest in zones where environmental heterogeneity and historical isolation intersect. These findings emphasize that GDI is not merely a refinement of existing diversity metrics but a complementary dimension that captures microevolutionary signals often overlooked in broad-scale biodiversity assessments.

### Climate stability predicts spatial patterns of GDI

GDI showed significant associations with both temperature and precipitation stability across Europe (all p ≪ 0.001). Correlation coefficients indicated moderate positive relationships, with stronger effects for temperature stability than precipitation. In linear models, temperature stability emerged as the dominant predictor of GDI (*β* = 0.30, *p* < 2 ×10^−16^), whereas precipitation stability had a weaker and slightly negative effect (*β* = −0.03, *p* < 10^−10^). Sampling intensity contributed only marginally (*β* = 0.01). Overall explanatory power was modest but significant (*R*^2^ ≈ 0.08) (Appendix Tab. S1). When accounting for spatial structure using GAMs, the smooth geographic term explained a large proportion of variance. Nevertheless, both climate stability components remained significant, and their effects became consistently positive, indicating that areas with greater long-term climatic persistence tend to harbor higher genetic diversity (Appendix Tab. S2). Effect size comparisons further emphasized the stronger contribution of temperature stability relative to precipitation stability (relative importance figure), and partial response curves confirmed monotonic increases of GDI with increasing stability (Fig. 5).

**Figure 5:**
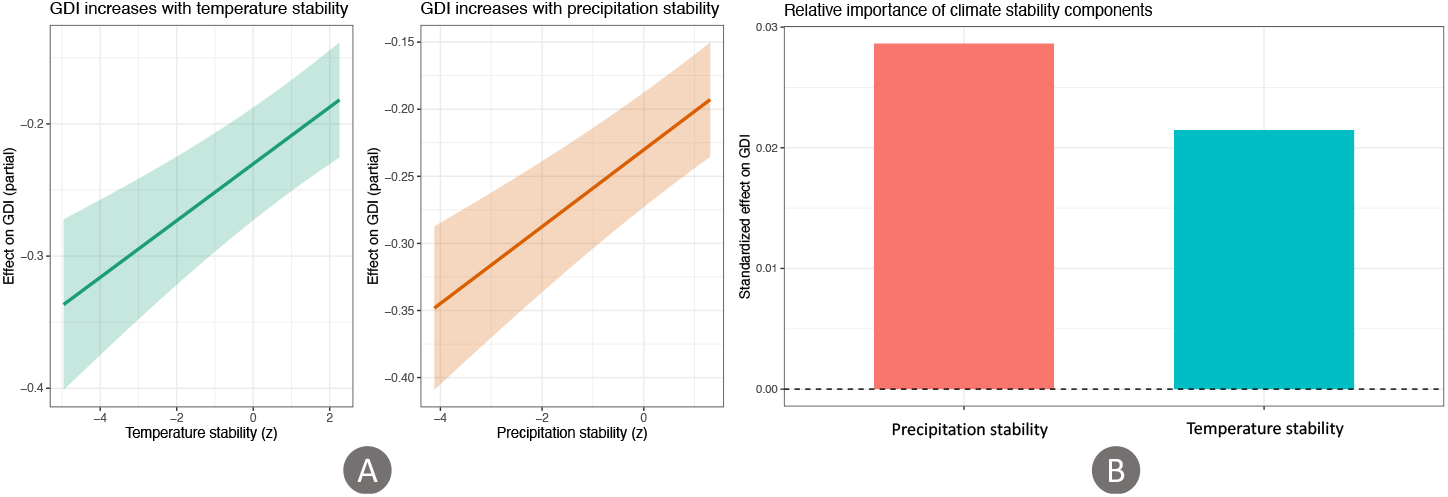
Associations between climate stability and genetic diversity. (**A**) Partial response curves from generalized additive models (GAMs) illustrating the relationship between GDI and each predictor while controlling for spatial structure. Shaded areas represent 95% confidence intervals. (**B**) Relative importance of predictors based on standardized coefficients from the linear model, showing that temperature stability has a substantially stronger influence on GDI than precipitation stability or sampling intensity.

## Discussion

In this study, we developed a composite Genetic Diversity Index (GDI) to integrate the genetic dimension of biodiversity into large-scale, multi-species assessments. Using European vascular plants as a case study, we demonstrated that millions of publicly available, geo-referenced genetic sequences can be effectively leveraged to identify hotspots of genetic diversity across broad spatial scales. Our approach highlights that GDI captures an emergent layer of biodiversity, rooted in evolutionary and demographic history, that is not fully reflected in taxonomic or phylogenetic diversity metrics. Recognizing and conserving such genetically rich regions is essential for maintaining evolutionary potential, especially under accelerating global change. These findings underscore the critical value of incorporating macro-scale genetic diversity assessments into biodiversity monitoring frameworks and conservation planning strategies.

### Implications for Biodiversity Conservation: how genetic diversity patterns relate to ecological and evolutionary processes

Spatial patterns of genetic diversity often reveal the imprints of both historical evolutionary processes, such as postglacial colonization, range shifts, demographic bottlenecks, and ongoing ecological dynamics, including environmental gradients, habitat heterogeneity, and species interactions (Hewitt, 2000; Hampe and Petit, 2005). For example, regions that served as glacial refugia during the Pleistocene tend to retain higher levels of intraspecific genetic diversity, as they hosted longterm stable populations with limited extinction-recolonization events (Petit et al., 2003; Provan and Bennett, 2008). In contrast, areas recolonized after glaciations typically show lower genetic diversity due to founder effects and reduced historical effective population sizes (Excoffier et al., 2009).

Climate stability is considered a major driver of spatial genetic diversity patterns and a proxy for long-term habitat suitability, as regions that experienced persistent conditions are more likely to have supported demographically continuous populations capable of retaining and accumulating genetic variation over time (Tzedakis et al., 2013; Hewitt, 2000; Hampe and Petit, 2005; Provan and Bennett, 2008). Importantly, the spatial signal inferred from refugial history is supported by our quantitative assessment of late-Quaternary climate stability. Across Europe, GDI increases significantly with the persistence of temperature regimes through time, even after accounting for spatial structure and sampling intensity (Fig. 5). This concordance indicates that regions experiencing reduced climatic fluctuation have tended to maintain larger or more continuous populations, thereby facilitating the long-term retention and accumulation of genetic variation. Although climate stability alone cannot explain all spatial heterogeneity in GDI, its consistent association across taxa suggests that macroevolutionary and demographic processes operating over glacial–interglacial cycles have left a detectable imprint on contemporary patterns of genetic diversity.

In our assessment of vascular plant genetic diversity, this becomes clear in regions such as Anatolia, European Alps and Carpathian, and Southern Iberian Peninsula, which stand out as the hotspots of genetic diversity (Fig. 3). These regions have collectively not only served as glacial refugia during the Pleistocene, allowing species survival and subsequent recolonization of Europe but also acted as corridors for post-glacial expansions, supporting a mix of relict and newly adapted plant populations (Feliner, 2011). On the other hand, low genetic diversity regions such as Scandinavia, Baltic region, Western Russia and British Isles experienced extensive glaciation during the Last Ice Age, leading to widespread population bottlenecks and reduced genetic diversity. The recolonization of these areas from limited post-glacial sources has further restricted genetic diversity (e.g.,Tollefsrud et al., 2008).

Ecologically, genetic diversity often correlates with environmental heterogeneity, as diverse habitats can promote population structuring, local adaptation, and long-term persistence of divergent lineages (Stein et al., 2014; Sexton et al., 2009). High habitat complexity may also buffer populations against demographic fluctuations, thus preserving genetic variation. A prime example of such environmental heterogeneity can be seen in the Alps and Dinaric Mountain systems which provide strong climatic and topographic gradients, fostering habitat specialization and genetic divergence (Hewitt, 2000; Petit et al., 2003). Similarly, in the regions of Southern Italy and Northern Tunisia, like other Mediterranean refugia, climatic stability and varied topography has contributed to the maintenance of diverse plant populations (Fig. 3). The presence of volcanic and coastal ecosystems further contributes to local adaptation and genetic differentiation (Médail and Diadema, 2009).

At a broader scale, ecotones and transition zones (e.g., between biomes or climate regimes) are frequently identified as hotspots of genetic diversity, where lineages of different evolutionary origins come into contact or hybridize (Kark, 2017; Zhou et al., 2024). Similarly, mountain systems and islands often harbor elevated genetic variation due to geographic isolation and opportunities for diversification (Gillespie and Roderick, 2002). The combination of mountains, drylands, and coastal ecosystems in Southern Iberian Peninsula has facilitated habitat differentiation, supporting diverse genetic lineages (Médail and Diadema, 2009). The Strait of Gibraltar serves as a biogeographic corridor, enhancing gene flow between Europe and North Africa (Rodríguez-Sánchez et al., 2008). Arguably, Anatolia’s varied topography and Mediterranean climate create numerous ecological niches, supporting high plant endemism and genetic diversity (Noroozi et al., 2021). In contrast, unlike the mountainous refugia in southern Europe, areas such as Parts of Central Eastern Europe (Poland, Czechia, Ukraine) and Western Russia lack complex topographies, resulting in lower habitat heterogeneity and fewer isolated populations, resulting in low levels of diversity (Feurdean et al., 2018).

By analyzing genetic diversity across multiple co-occurring species, it becomes possible to detect emergent patterns that transcend species-specific histories and reflect shared ecological or biogeographic drivers (Miraldo et al., 2016; Theodoridis et al., 2020). Such community-level genetic diversity metrics can reveal regions where evolutionary and ecological processes converge to shape biodiversity, providing insights into the mechanisms maintaining genetic variation at broader scales. This integrative perspective is crucial for understanding how genetic diversity is distributed across landscapes and how it may respond to future environmental changes.

### Potential applications of the genetic diversity index in conservation planning and biodiversity monitoring

The Genetic Diversity Index (GDI) developed in this study provides a scalable and integrative measure of intraspecific genetic diversity across multiple species and regions. Unlike traditional measures of biodiversity that focus on species presence or evolutionary lineage depth, GDI reflects population-level variation, which is crucial for long-term evolutionary resilience and adaptability. Here, we outline three key ways in which GDI can contribute to conservation planning, from global to local scales.

#### i. Revealing Genetic Hotspots Beyond Species Richness and Endemism

Numerous studies have shown that areas rich in species (taxonomic diversity) or phylogenetic lineages do not always coincide with areas of high intraspecific genetic diversity (Afonso Silva et al., 2025; Kahilainen et al., 2014). Our analysis, which combines taxonomic, phylogenetic, and genetic diversity patterns of vascular plants, confirms that relying solely on species richness or endemism may miss regions of critical evolutionary value. This is evident in our comparison of currently known plant diversity hotspots of Europe with patterns of GDI (Fig. 3). For example, glacial refugia such as the Iberian Peninsula, the Balkans, and parts of southern Italy have been repeatedly identified as hotspots of genetic diversity, even when species richness is not exceptionally high (Fig. 3) (Taberlet et al., 2012; Médail and Diadema, 2009). These regions often harbor deep intraspecific lineages that reflect long-term population persistence during climatic oscillations.

By incorporating GDI into hotspot identification frameworks (e.g., Myers et al., 2000; Daru, 2024; Macher et al., 2024), conservation planners can recognize regions with high evolutionary potential, areas that not only harbor many species, but also the genetic depth necessary for future adaptation and speciation. GDI thus adds an essential dimension for prioritizing regions under threat from climate change, land use, and fragmentation. Further, GDI can help identify cryptic refugia or microhabitats that maintain unique genetic variants, which may be overlooked in traditional assessments focused on species counts alone (Hewitt, 2000; Provan and Bennett, 2008).

#### ii. Taxon-Specific Conservation Planning and Functional Insights

The GDI can also be applied to conservation planning focused on specific taxonomic groups, such as forest trees, pollinators, or crop wild relatives. For example, studies on European oaks, spruce, and beech have shown that genetic diversity patterns are shaped by complex postglacial recolonization dynamics and long-term ecological processes (Magri et al., 2006; Kremer and Hipp, 2020; Karunarathne et al., 2024). In such taxa, genetic diversity is often heterogeneously distributed across their range, reflecting local adaptation, demographic history, and gene flow barriers.

When sampling within a given area or across a taxonomic group is evenly distributed, using the mean GDI across species (or populations) may obscure underlying spatial patterns of genetic diversity. This is because mean GDI flattens the heterogeneity inherent to individual species’ diversity distributions, often resulting in uniform or uninformative surfaces. In contrast, summing the GDI (*s*GDI) across all sampled taxa amplifies shared diversity signals and reveals emergent patterns, such as biodiversity hotspots driven by overlapping centers of intraspecific variation. For example, in our analysis, while mean GDI showed little spatial structure due to the variability among individual species, the cumulative index (*s*GDI) revealed a pronounced central hotspot (Fig. 6). This illustrates how *s*GDI better captures the collective functional and evolutionary value of a community, making it ideal for conservation approaches targeting multi-species genetic resilience.

**Figure 6:**
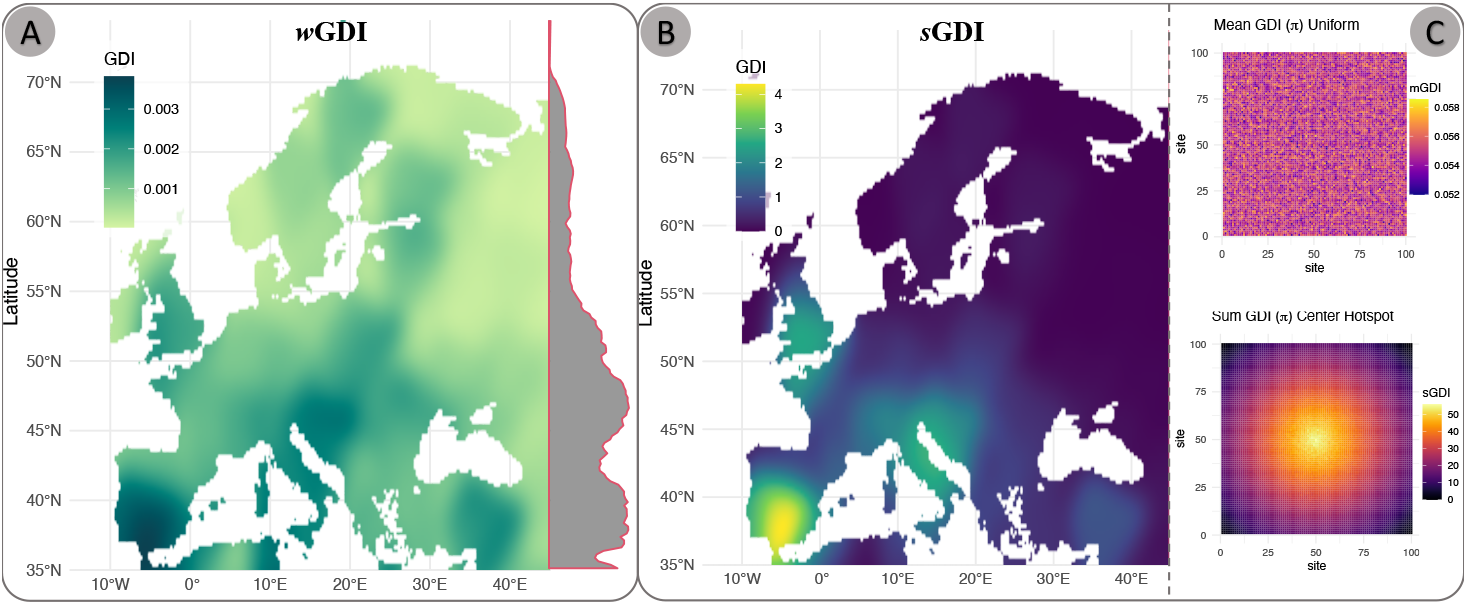
Two alternative genetic diversity indices for different scenarios of genetic diversity assessments based on sampling and species richness. **A**. Cumulative GDI (*s*GD I; sum of *π* across all species at a site, *sGDI* = Σ*π*_*i*_). **B**. Weighted GDI (*w*GDI; mean *π* weighted by species richness, *wGDI* = (Σ*π*_*i*_/*S*) × (*S*/*S*_total_)^α^). **C**. A hypothetical example of the cumulative genetic diversity index (*s*GDI) in an ideal scenario where all species are equally sampled across a grid cell. The *s*GDI is calculated as the sum of nucleotide diversity across all species in the grid cell, providing a comprehensive measure of genetic diversity. This approach allows for a more nuanced understanding of genetic variation and its implications for conservation planning and biodiversity monitoring, whereas the mean nucleotide diversity (*m*GDI) may not fully capture the genetic richness of the area. Note: the same *π* matrix was used to calculate both *s*GDI and *m*GDI in the figure. Weighted GDI (*w*GDI) provides a more balanced view of genetic diversity across species. It is particularly useful in scenarios where species richness varies significantly across grid cells and the sampling is near complete, allowing for a more equitable representation of genetic diversity.

For groups like insects, which are often understudied in terms of genetic data, using a GDI framework can help identify regions where ecological interactions and microhabitat heterogeneity contribute to genetic structure (Kerr et al., 2015; Müller et al., 2018). In tropical tree species, genetic diversity has been linked to both historical stability and dispersal syndromes, highlighting the value of integrating GDI into tropical forest conservation planning (Hardy et al., 2013). By layering taxonomic diversity with GDI, conservation plans can move beyond presence/absence data and incorporate evolutionary and ecological depth, enabling strategies that promote long-term

#### iii. Fine-Scale Applications: Islands, Fragmented Habitats, and National Assessments

At smaller spatial scales, the GDI offers valuable insights for island biogeography, habitat fragmentation studies, and national biodiversity assessments. For instance, islands such as Crete or Corsica may exhibit relatively low species richness but disproportionately high genetic endemism due to isolation and long-term lineage divergence (Panitsa et al., 2006; Lindenmayer, 2019), a pattern consistent with the elevated GDI scores observed in our results for other topographically or historically isolated regions (see Fig. 3A & 6B). Similarly, in fragmented habitats such as alpine meadows or Mediterranean woodlands, local populations may harbor unique allelic combinations that contribute disproportionately to regional adaptive potential, even when overall species richness is low.

In biodiversity hotspots where habitat fragmentation and human activity have intensified (e.g., the Atlantic Forest of Brazil, Southeast Asian tropical forests, and the Congo Basin), the weighted Genetic Diversity Index (*w*GDI) becomes particularly informative: by accounting for species richness, it can identify sites where much of the standing genetic diversity is concentrated in a few surviving species (see Fig. 6B), thereby prioritizing areas at greatest risk of genetic erosion. In this way, both GDI and its derivatives offer scalable, fine-resolution tools that can guide conservation strategies across island systems, fragmented landscapes, and national biodiversity planning efforts.

#### iii. Fine-Scale Applications: Islands, Fragmented Habitats, and National Assessments

At smaller spatial scales, the GDI provides valuable insights for island biogeography, habitat fragmentation studies, and national biodiversity assessments. For instance, Mediterranean islands such as Crete and Corsica, though relatively species-poor, exhibit disproportionately high levels of genetic endemism due to isolation and long-term lineage divergence (Panitsa et al., 2006; Lindenmayer, 2019). This aligns with our findings of elevated GDI, particularly *w*GDI, in historically and topographically isolated regions (see Fig.3A and Fig.6B).

In fragmented landscapes such as alpine meadows or Mediterranean woodlands, local populations can harbor unique allelic combinations that disproportionately contribute to regional adaptive potential, even when species richness is low. In such contexts, a GDI adjusted only for sampling intensity may obscure these conservation-relevant patterns by assigning equal weight to all species, regardless of their differing contributions to genetic diversity. This is where the weighted Genetic Diversity Index (*w*GDI) is especially useful.

By factoring in species richness, *w*GDI highlights regions where a large fraction of standing genetic diversity is concentrated in just a few species. In our analysis, this improved resolution allowed the identification of priority areas—including Mediterranean islands—where persistent habitat fragmentation and species loss make remaining genetic diversity particularly vulnerable (see Fig. 6B). More broadly, although beyond the scope of our current study, this weighted approach would be well suited to other biodiversity hotspots experiencing similar pressures, such as the Atlantic Forest, Southeast Asian archipelagos, or the Congo Basin.

The GDI framework can also help identify remnant habitat patches that retain high intraspecific diversity, offering a practical tool for restoration planning, connectivity design, and the conservation of genetic resources. In temperate regions like European lowland grasslands and Mediterranean scrublands—where fragmentation is driven by agriculture and urbanization—the GDI enables the detection of evolutionarily significant areas that might otherwise be overlooked in traditional species-focused assessments.

In national conservation frameworks such as Red List evaluations, biodiversity strategies, or Natura 2000 site planning (Bundesamt für Naturschutz (BfN), 2025), GDI could be used to flag genetically important regions that do not qualify under conventional criteria. This aligns with the goals of the Post-2020 Global Biodiversity Framework, which calls for the integration of genetic diversity into conservation monitoring and policy (CBD, 2020). By adopting GDI and its derivatives, such initiatives can more effectively target areas where genetic resources are most at risk, improve landscape connectivity, and better quantify the evolutionary impacts of conservation efforts.

### Limitations and Data Gaps in Multi-Species Genetic Diversity Assessment

Our meta-analysis reveals uneven availability of geo-referenced genetic data across European plant families. Several species-rich and ecologically dominant groups, including Asteraceae, Rosaceae, and Ranunculaceae, remain underrepresented, whereas some small families are disproportionately sampled (Appendix Fig. S6–S7). These disparities reflect limitations of current data resources rather than the GDI framework itself, yet they inevitably influence how completely regional patterns of intraspecific variation can be recovered.

Missing key ecological contributors may reduce the ability of GDI to capture functional and historical signals. Dominant species are known to structure ecosystems and shape landscape-level genetic patterns (Ellison et al., 2005; Isbell et al., 2011), whereas localized overrepresentation of narrow-ranged taxa may inflate apparent regional diversity. Recognizing such imbalances therefore provides a roadmap for targeted improvement and more representative sampling.

An important avenue for development is the inclusion of ecological weighting based on rarity, abundance, or functional importance. Rare species may preserve unique evolutionary lineages (Rodriguez-Gacio et al., 2009), while widespread species with large, structured populations contribute strongly to macroecological patterns of adaptive potential. Incorporating these dimensions would allow GDI to better reflect the unequal contributions of species within assemblages.

Despite these challenges, GDI yields robust and interpretable spatial patterns where (i) local species richness is high and (ii) a sufficient fraction of taxa has been genetically characterized (Fig. 4A). Areas with sparse taxonomic representation, particularly in northern and eastern Europe, show greater uncertainty (Fig. 4B), mirroring a general constraint in biodiversity data rather than a methodological artifact.

These observations are consistent with previous macrogenetic syntheses (Miraldo et al., 2016; Theodoridis et al., 2020) and emphasize the value of expanding sequencing coverage, prioritizing both foundation and rare taxa, and integrating ecological attributes into future iterations of the index. Such developments will further increase the accuracy, representativeness, and conservation relevance of large-scale genetic diversity assessments.

## Future Directions: Toward Scalable and Integrative Genetic Diversity Monitoring

Rapid growth in whole-genome sequencing (WGS) and genotyping-by-sequencing (GBS) is transforming the potential for macrogenetic inference. While the present framework relies on marker-based nucleotide diversity (*π*), future developments can incorporate genome-wide information to estimate complementary metrics such as heterozygosity (*He, Ho*), inbreeding coefficients (*F*_*IS*_), and population differentiation (*D*_*xy*_, *F*_*ST*_, Bayesian ω; Gautier, 2015). Although broad application of such data remains constrained by computational demands and uneven curation, they are already enabling deeper analyses in priority clades including crops, trees, and narrow-range endemics (Parchman et al., 2018; Luikart et al., 2019).

Importantly, GDI currently quantifies neutral genetic diversity, providing a proxy for demographic stability and long-term evolutionary potential. It therefore offers a scalable baseline upon which adaptive dimensions can be layered. Advances in landscape genomics and genotype–environment association approaches increasingly allow the detection of loci associated with climate tolerance and local adaptation (e.g., Fitzpatrick and Keller, 2015; Rellstab et al., 2015). Integrating these signals with GDI will make it possible to distinguish regions that harbor not only large reservoirs of standing variation but also alleles relevant for future environmental change, thereby supporting climate-smart conservation planning.

The effectiveness of this framework depends on georeferenced genetic data, which are expanding rapidly but remain unevenly distributed across taxa and regions. Rather than limiting inference, these gaps define clear priorities for innovation. Emerging artificial intelligence (AI) and machine-learning methods can assist in imputing missing metadata, detecting sampling biases, predicting diversity patterns, and guiding targeted data acquisition (e.g., Yoosefzadeh-Najafabadi et al., 2024; Daru, 2024). Coupling AI-assisted harmonization with automated, reproducible pipelines will substantially increase scalability and robustness, even where empirical coverage is incomplete.

Although developed for European vascular plants, the framework is inherently global. A preliminary analysis using 1 million georeferenced accessions demonstrates strong representation in Europe, North America, Southeast Asia, and Australia, but also reveals persistent shortfalls in South America, Africa, Western Asia, and Siberia. These disparities largely reflect research infrastructure rather than biological relevance. Encouragingly, simulations show that reliable inference is achievable once taxon-specific data thresholds are reached, indicating that strategic sampling can compensate for incomplete coverage.

Achieving a global implementation will require coordinated international collaboration, standardized metadata practices, and integration of datasets currently held within regional institutions. Computational challenges associated with large-scale alignment, modeling, and synthesis are substantial but tractable using existing high-performance and cloud-based infrastructures.

Taken together, continued improvements in data availability, harmonization, and analytical integration position the GDI framework to become a central component of next-generation biodiversity monitoring. By linking neutral diversity, adaptive potential, and spatial prediction within a unified architecture, it offers a pathway toward globally consistent identification of evolutionary resources for conservation.

## Data, Scripts and Software Accessibility

Species lists, genetic meta-data tables, species distribution tables, sequence alignments, individual nucleotide diversity matrices, and the GDI outputs along with all the scripts and code used to generate the GDI and the figures are available on Zenodo (https://doi.org/10.5281/zenodo.18632565). Functions used in the analysis are also available in the R package gSoup (https://doi.org/10.5281/zenodo.15624400).

## Author Contributions

P.K. conceived the study; M.K. compiled and processed the data; P.K. and M.K. developed the GDI framework, performed analyses, and generated figures; P.K., M.K., and L.E.R. interpreted results; P.K. drafted the manuscript with input from M.K. and L.E.R.; all authors revised and approved the final version.

