## appendices for "Revealing Evolutionary Signals Across Landscapes: A Scalable Genetic Diversity Index for Multi-Species Analysis"

#### Appendix 1: COMPLETE WORKFLOW

The overall workflow was structured into two main components:

1. Data acquisition and meta-data analysis, which involved downloading, filtering, and processing of species occurrence records, metadata of genetic data, and DNA sequence information for European vascular plants; and

2. Diversity analysis and mapping, where species richness, phylogenetic diversity, and geographic patterns of genetic diversity were quantified and visualized

An abstract overview of the key analytical steps is presented in main text Fig. 1A and 1B.

##### Data acquisition and meta-data analysis

###### Occurrence data

To compile a comprehensive list of native vascular plant species occurring in Europe, we used the Plants of the World Online database (POWO; `https://powo.science.kew.org/`) and the World Checklist of Vascular Plants (WCVP; Govaerts et al., 2021). The WCVP was used as the source to extract metadata for all available nucleotide sequences (accessions) for each species. The complete dataset from WCVP was refined to include only currently accepted species that are native to Europe. This filtering process involved the following criteria: i) taxonomically accepted species, ii) native status, iii) exclusion of subspecies, iv) extant species only, and v) geographical occurrence limited to Europe following EEA biogeographic regions (European Environment Agency, 2025) . The resulting plant list of 26,242 species (included in the GitHub repository) formed the foundation for all subsequent analyses.

###### Species Distributions

We obtained species distribution data from two sources: a) 11,276 species were obtained from Daru (2024) dataset, whose method models a species' realized niche by incorporating: i) spatial probability of recorded occurrences, ii) dispersal potential estimated via a diffusion coefficient (D), and iii) the natural distribution range of the species' respective plant family; b) for species not included in this dataset, yet we obtained genetic data for (1,298 species), we applied the same modeling approach as Daru (2024) to estimate their geographic distributions. It is a standardized workflow based on cleaned occurrence data and environmental predictors. Occurrence records were downloaded from GBIF and from the genetic metadata compiled in this study, and taxonomically harmonized using the World Checklist of Vascular Plants (WCVP). Records were spatially cleaned with the `CoordinateCleaner` (Zizka et al., 2019) package to remove duplicates, erroneous coordinates, and cultivated specimens. For each species with at least five unique localities, alpha hull polygons were generated using the `rangeBuilder` (Davis Rabosky et al., 2016) package to approximate native ranges. Environmental predictors (19 bioclimatic variables from WorldClim v2.1 and elevation: Fick and Hijmans (2017)) were reduced for collinearity using variance inflation factor (VIF) filtering. Species distribution models were implemented using MaxEnt under a 5-fold cross-validation framework via the `phyloregion` (Daru et al., 2020) package, with optimized feature classes and regularization multipliers. Model performance was evaluated using AUC, TSS, and Boyce Index, and continuous suitability outputs were converted to binary presence–absence maps using the equal sensitivity–specificity threshold. Final species range maps were rasterized at 0.25° resolution for further diversity analyses.

#### Genetic meta-data

Metadata for nucleotide sequences of vascular plant species native to Europe were retrieved from the NCBI GenBank database using a custom script built with the R package rentrez (Winter, 2017), which interfaces with NCBI's E-utilities API (Sayers et al., 2011). For each species on the refined plant list (26,242 species), we retrieved metadata associated with all available sequence accessions. Specifically, the extracted metadata included: i) sequence definition (containing marker and species details), ii) sequence ID, iii) locus, iv) publication information, v) author(s), vi) collector(s), vii) collection date, viii) collection location, ix) geographic coordinates, x) accession number, xi) species name, xii) submission date, and xiii) sequence length.

#### Filtering

Using georeferenced information (i.e., geographic coordinates and collection locations), we retained only those accessions suitable for spatial genetic diversity analysis. These accessions were classified into two categories based on the type of georeferencing: i) primary georeferenced accessions: records with precise geographic coordinates, and ii) secondary georeferenced accessions: records with descriptive location information (e.g., country, city, locality). Secondary georeferenced accessions were geocoded (i.e., converted into coordinates) using the open-source R packages tmaptools (Tennekes, 2018) and tidygeocoder (Cambon et al., 2021). Accessions containing only vague or unresolvable location data (e.g., only a country name) were excluded from the spatial genetic diversity analysis but were retained for data availability assessments.

#### Marker assignment

To identify commonly used DNA markers across European vascular plant species, we scanned over 25 million GenBank accession definitions linked to taxa native to Europe. The objective was to extract marker-level information for downstream genetic diversity analyses.

Many markers were inconsistently labeled in GenBank entries (e.g., "matK" vs. "maturase K", or "ITS" vs. "internal transcribed spacer 1/2"). To address this, we compiled a comprehensive list of synonymous terms for each marker and used them to search the definition fields. Once detected, accessions were assigned to a standardized marker label to avoid duplication and ensure consistency. We also screened for whole genome sequences (WGS). However, these were excluded from the genetic diversity analyses to minimize biases introduced by sequence incompleteness and inconsistent representation across taxa. All custom R scripts used to perform marker extraction and assignment are available in the Zenodo repository (`https://doi.org/10.5281/zenodo.18632565`).

Because genetic markers were not consistently annotated in the sequence metadata retrieved from GenBank, we implemented a custom script to infer marker identity directly from the definition fields of each accession. The script scanned for more than 50 (see list below) commonly used plant markers and barcodes (e.g., *matK*, *rbcL*, ITS) and assigned each sequence to its corresponding species–marker group (e.g., *Picea abies* – *matK*, *Quercus petraea* – ITS). To prevent redundancy, only one unique accession per species–marker combination was retained. Downstream analyses focused on species–marker sets with at least five georeferenced accessions.

**List of scanned markers:** matK, maturase K, rbcL, psbA-trnH, ITS, ITS1, ITS2, trnL-trnF, atpB-rbcL, COI, ycf1, trnK, ndhF, rpl16, accD, ndhJ, rpoB, rpoC1, psbB, psbC, atpF-atpH, petA-psbJ, trnS-trnG, petD, clpP, rps16, rps4, rpl20-rps12, rpl14, rbcL-atpB, psbM-trnD, psbB-psbH, trnS-rpS4, GBSSI, waxy, PHYC, LEAFY, ncpGS, G3pdh, Adh, CHS, F3H, cox1, cox2, cob, nad1, nad2, nad4, nad5, atp1, atp6, 18S rRNA, 26S rRNA, 28S rRNA, 5.8S rRNA, ETS, WGS (whole genome sequence).

##### Sequence download

FASTA sequences corresponding to the selected, georeferenced, marker-assigned accessions were downloaded from GenBank using the NCBI Entrez Direct (EDirect) command-line tools. Over 12 million sequence accessions were retrieved and stored in separate FASTA files, organized by species-marker combinations, for subsequent processing and diversity analyses. We applied several quality control steps during and after the download process to ensure data integrity, including: i) removing duplicate sequences, ii) filtering out sequences with ambiguous bases (e.g., N's), iii) excluding sequences below a minimum length threshold (specific to each marker), and iv) verifying taxonomic names against the WCVP to ensure consistency.

#### Diversity analyses

##### Species richness and phylogenetic diversity

Taxonomic diversity was calculated as the total number of vascular plant species present within each spatial unit, defined as a raster cell of at 0.25° raster cell resolution on the WGS 84 coordinate reference system (~ 20 km). Species presence/absence maps generated from species distribution models (SDMs) were overlaid, and the number of unique species per raster cell was computed. This resulted in a species richness map across Europe. Our analysis included approximately 50% of all vascular plant species occurring in the European region, representing a dataset of 12,571 species.

Phylogenetic diversity (PD) was assessed using Faith's Phylogenetic Diversity index (Faith, 1992), implemented via the phyloregion R package (Daru et al., 2020). We employed the comprehensive, dated vascular plant phylogeny developed by (Smith and Brown, 2018), which was pruned to retain only the species present in our compiled presence/absence matrix. Two input matrices were used to compute PD: i) a community matrix representing the presence or absence of each species within raster cells, and ii) a phylogenetic distance matrix including branch lengths among the included species. Following the methodology outlined in Daru et al. (2020), we further calculated a weighted version of Faith's PD index, correcting for differences in species richness across sites by standardizing the effect size. This produced a weighted PD community matrix, which was subsequently merged with the rasterized European grid (0.25° raster cell) for spatial visualization and analysis.

##### Spatial Genetic Diversity & Genetic Diverstity Index (GDI) Development

Geographic genetic diversity was assessed by calculating the mean nucleotide diversity ($\pi$) of all individuals of all species occurring within a given spatial unit (0.25° raster cell). Nucleotide diversity ($\pi$) quantifies genetic variation by measuring the average number of nucleotide differences per site between two randomly chosen sequences (Nei and Li, 1979), and is calculated as:

$$\pi = \sum_{ij} x_i x_j k_{ij}$$

Where $x_i$ and $x_j$ are the frequencies of the ith and jth sequences in the population, and $k_{ij}$ is the number of nucleotide differences per site between them.

##### Per-Individual and Per Grid Cell (per-site) $\pi$ Calculation:

To estimate nucleotide diversity at the individual level, we computed the average pairwise nucleotide diversity ($\pi_i$) across all sequences for each species–marker combination. When multiple markers were available for a species, we retained the marker with the highest number of sequences to maximize sample size and geographic coverage, while also making sure that only homologous sequences were aligned and compared. Multiple sequence alignments were performed using MAFFT (Katoh and Standley, 2013), with the reverse complement option and settings optimized for maximum accuracy. All alignments were manually inspected to ensure proper alignment quality and to correct any misalignments; We applied a threshold of 80% sequence coverage to filter out poorly aligned or incomplete sequences. Further, species-marker combinations whose sequences that were entirely conserved across all individuals (i.e., all pair-wise $\pi = 0$) were excluded from further analysis, as they do not contribute to genetic diversity estimates. For each sequence record, the resulting mean $\pi$ value was assigned and spatially linked to its geographic origin. These values were mapped onto a raster grid at 0.25° resolution on a WGS84 map projection. To reduce sampling bias from overrepresented sites, we applied a subsampling approach, averaging genetic diversity across all samples within each site.

We used these per-individual nucleotide diversity ($\pi$) to calculate summary statistics for different taxonomic levels (e.g., species, genus, family). For each species, we computed the mean nucleotide diversity ($\pi$) across all individuals sampled within its modeled native range (based on species distribution maps). This provided a species-level estimate of genetic diversity that accounts for geographic variation within the species' distribution. These summaries show the distribution of genetic diversity across taxonomic groups and allowed us to identify patterns of variation at different hierarchical levels (e.g., see main text section 'Genetic Diversity Patterns in Eurpean Vascular Plants' and Appendix Fig. S1).

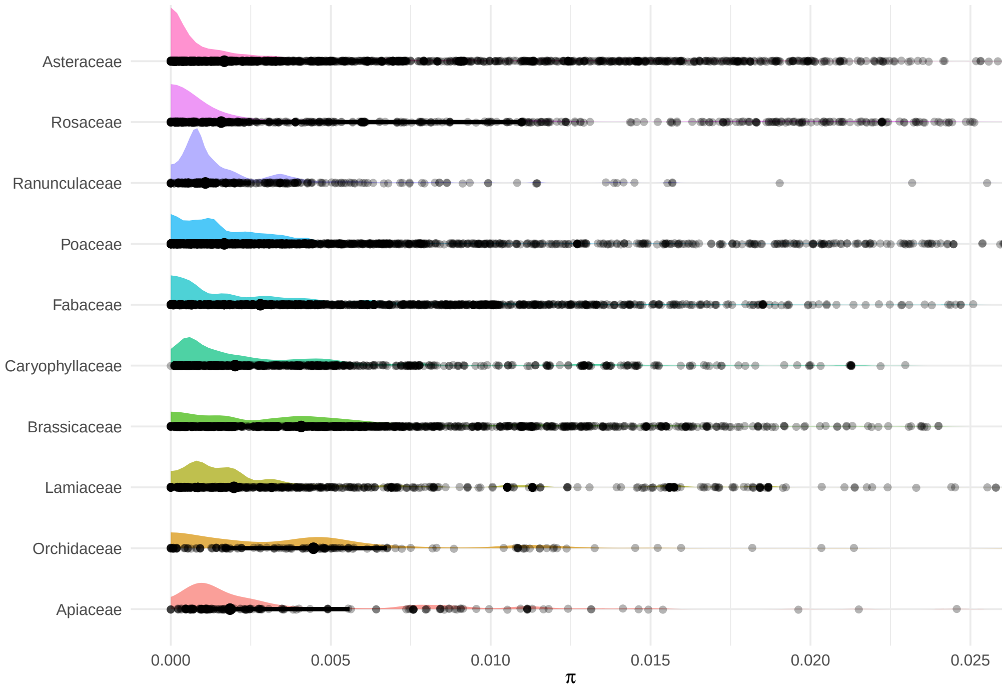


Figure S1: Nucleotide diversity variation among top 10 most specious families in Europe. The curves show the Kernel density distribution of $\pi$ values of all the species in each family. Note: the x-axis has been trimmed at 0.025 to improve the visibility of the curves.

#### Spatial Estimation of Genetic Diversity within Species Ranges:

To avoid complete omission of under-sampled regions while preserving biological realism, we applied a conservative spatial interpolation procedure to estimate nucleotide diversity ($\pi$) within each species' modeled native range (based on SDM output). Rather than extrapolating across broad geographic gaps, $\pi$ values were interpolated only between empirically sampled locations located within a maximum geographic distance threshold and separated by no known topographical or ecological barrier (e.g., mountain ranges, major water bodies; see Appendix 4). Estimates were calculated using inverse-distance weighting (IDW) based on the nearest sampled populations (default $k = 3$), and only propagated into unsampled grid cells with direct spatial connectivity to sampled neighbors. No values were assigned outside SDM-predicted distributions or across known barriers to gene flow.

This conservative interpolation approach avoids smoothing across genetic discontinuities and functions as a gap-filling tool rather than a predictor of unobserved diversity. All interpolated values are flagged and accompanied by a confidence score based on sampling density and neighborhood structure (see main Fig. 3B and Methods). The interpolation function and default constraints are implemented in the `wind_interpolate()` routine in the R package `gSoup` (Karunarathne, 2024). We emphasize that interpolated $\pi$ values serve as spatial placeholders in sparsely sampled regions and are not interpreted as evidence of true genetic continuity; rather, they allow regional-scale diversity estimates while explicitly highlighting areas of insufficient empirical sampling.

**Multi-Species Genetic Diversity Index (GDI):**

To characterize patterns of multi-species genetic diversity across Europe, we overlaid the spatially extrapolated nucleotide diversity ($\pi$) maps of all species. From this composite, we computed three complementary Genetic Diversity Indices (GDIs) to quantify collective genetic diversity at each site. Each index captures a different aspect of spatial genetic variation catered for different sampling and study purposes, and provides unique biological insights, as detailed below.

- Mean Genetic Diversity Index (mGDI):

  The mean genetic diversity (mGDI) was calculated as the average $\pi$ value of all sampled species at a site:

$$mGDI = \frac{1}{S}\sum_{i=1}^{S}\pi_i$$

  where $S$ is the number of species sampled at a site, and $\pi_i$ is the nucleotide diversity of each species.

  mGDI reflects the average within-species genetic diversity for the sampled community, independent of species richness. This allows comparisons of genetic variation per species, enabling identification of areas where species tend to be more genetically diverse on average, regardless of how many are present. It is more robust to sampling intensity than cumulative GDI (sGDI; see below) but may downplay the influence of particularly diverse taxa in rich communities; hence we call it 'uncorrected GDI' in the main text. We used this index for initial exploratory analyses of genetic diversity patterns across Europe (see Appendix Fig. S2).

- Corrected Genetic Diversity Indices (cGDI):

  To mitigate potential sampling biases, such as uneven species representation or under-sampling, we implemented three correction methods to the mean genetic diversity index (mGDI). These corrected indices standardize or adjust diversity estimates and enable more accurate comparisons across sites with varying levels of taxonomic coverage. The three correction methods are:

  1. Regression-Based Correction (cGDI):

We fit a linear model to predict GDI as a function of sampling proportion at each site:

$$GDI = \beta_0 + \beta_1 \left( \frac{S}{S_{\text{total}}} \right)$$

Where GDI is the observed genetic diversity index at a site, $S$ is the number of sampled species, $S_{total}$ is the total known species richness (from species distribution maps), $\beta_0$ is the intercept, $\beta_1$ is the slope coefficient.

The corrected GDI (cGDI) was then calculated using the model residuals:

$$cGDI = GDI - \beta_1 \left( \frac{S}{S_{\text{total}}} \right)$$

cGDI removes the linear dependency between observed genetic diversity and sampling proportion. This approach isolates the true biological signal of genetic diversity by statistically controlling for sampling bias. It is especially powerful when sampling effort correlates with geographic or ecological gradients. We used this index for the main spatial analyses presented in the manuscript.

2. Weighted by Species Richness (wGDI):

   We adjusted diversity values by weighting them relative to total species richness at a site:

$$wGDI = \left( \frac{\sum_{i=1}^{S} \pi_i}{S} \right) \times \left( \frac{S}{S_{total}} \right)^{\alpha}$$

   where $\pi_i$ is the nucleotide diversity of the ith sampled species, $S$ is the number of sampled species, $S_{total}$ is the known total species richness for the site, and $\alpha$ is a scaling exponent that moderates the influence of low-sampling bias.

   The wGDI adjusts for incomplete taxonomic sampling by factoring in how much of the local flora was actually sampled. This helps ensure that sites with low sampling coverage are not over- or underrepresented in the final diversity assessment. The exponent $\alpha$ can be tuned emperically based on confidence in sampling completeness as follows:

   *Empirical calibration of alpha:* The scaling factor ($\alpha$) controls the strength of the correction for incomplete sampling. We empirically estimated $\alpha$ by fitting the relationship between mean nucleotide diversity and sampling completeness ($S/S_{total}$) across well-sampled grid cells, using the slope of a log–log regression. In sensitivity analyses, we also evaluated fixed values ($\alpha$ = 0.5, 1, 2) to test the robustness of spatial patterns. We explain the ideal use of this index in the discussion in the main text.

3. Sample-Based Rarefaction (rGDI):

   We implemented a rarefaction approach by randomly subsampling 20% of the sampled species at each site. This subsampling was repeated 10,000 times, and the mean diversity value was computed across iterations to produce the rarefied GDI.

   The rGDI offers a sampling effort-independent estimate of average genetic diversity by standardizing diversity estimates across different sample sizes. It is particularly valuable when sampling effort is highly uneven, allowing fairer comparisons between well- and poorly-sampled sites. It also requires a high intensity of sampling for better accuracy.

These corrected GDIs were normalized to range from 0 to 1, where 0 indicates no observed genetic diversity or no species sampled at a site, and 1 represents the maximum observed genetic diversity, corresponding to full species sampling and all species exhibiting the highest possible nucleotide diversity ($\pi = 1$). However, note that a value of $\pi = 1$ typically indicates alignment artifacts or comparisons between highly divergent sequences, such as different species.

These indices inherently adjust for sampling completeness, making them more suitable for comparative spatial analyses of genetic diversity across landscapes. If only a subset of species is sampled ($S < S_{total}$), the index reflects both the observed genetic variation and the extent of species coverage.

- Cumulative Genetic Diversity Index (sGDI):

  The cumulative genetic diversity (sGDI) was calculated by summing the $\pi$ values of all species occurring at a site:

  $$sGDI = \sum_{i=1}^{S} \pi_i$$

  where $\pi_i$ is the nucleotide diversity of the $i$th species at a given site, and $S$ is the number of sampled species at that site.

  The sGDI captures the total genetic variation present at a site across all sampled species. This metric emphasizes cumulative diversity, making it useful for identifying hotspots of overall genetic richness. However, it is sensitive to species richness and sampling intensity, sites with more species or better sampling will naturally have higher values. Our Discussion section explains the benefits and ideal cases where this index is useful.

**Assumptions and Considerations:** i) equal weighting - Each species contributes equally to the index, irrespective of its ecological role or abundance. While this simplifies calculations, functional or ecological weighting schemes could be incorporated in future extensions, ii) sampling representation - though representative sampling enhances accuracy, the correction methods minimize bias introduced by uneven sampling intensity, offering a pragmatic and scalable solution for macroecological genetic assessments. Together, these GDIs provide a robust framework for quantifying and comparing genetic diversity across spatial scales and taxa, depending on the focus of the study.

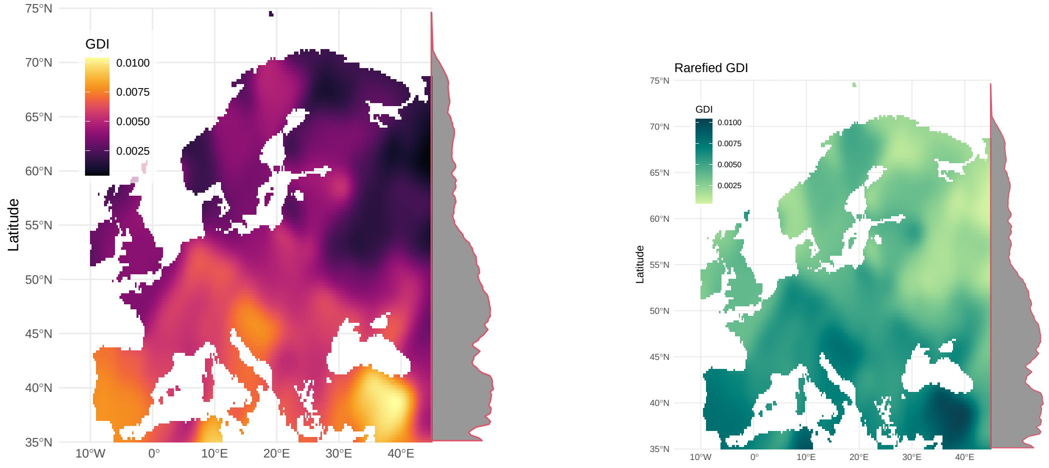


(A) mGDI (B) rGDI

Figure S2: Spatial distribution of uncorrected mean genetic diversity index (mGDI) (A) and rarefied genetic diversity index (rGDI) (B) of vascular plants across Europe . The mGDI represents the mean nucleotide diversity of all species with available sequence data occurring within each raster grid cell. For each cell, mGDI was calculated by averaging the uncorrected nucleotide diversity ($\pi$) values across all species present. This metric reflects the average intraspecific genetic variation per site and provides a comparative measure of local genetic diversity across the European flora. Regions with higher mGDI values indicate sites where species tend to exhibit higher genetic variation. The rGDI was calculated by randomly subsampling 20% of the species at each site and averaging the resulting $\pi$ values across 10,000 iterations. This rarefaction approach standardizes genetic diversity estimates across sites with varying sampling intensity, providing a more comparable measure of genetic diversity that accounts for differences in species representation. The spatial patterns observed in rGDI highlight areas of high genetic diversity while minimizing bias from uneven sampling effort. Both maps are visualized using a color gradient where warmer colors indicate higher genetic diversity.

#### Validation of Bias Correction Methods

To evaluate the effectiveness of our correction methods in minimizing sampling bias, we conducted a series of validation analyses using the final multi-species GDI dataset.

**Correlation-Based Validation:** We first assessed the relationship between genetic diversity indices and potential sources of bias, particularly sampling intensity (i.e., the proportion of species sampled per site). Specifically, we examined: i) the correlation between uncorrected GDI values and the proportion of species sampled at each site (i.e., sampling intensity), and ii) the relationship between corrected and uncorrected GDI values across sites with varying species richness.

Our results showed no strong or consistent trends between corrected GDI and sampling proportion (Pearson's r = –0.0001), indicating that the correction methods successfully reduced bias due to uneven sampling. Additionally, comparison plots between corrected and uncorrected GDI values revealed non-linear convergence at low richness sites, further supporting the robustness of our approach.

**Permutation Tests:** To test the sensitivity of GDI estimates to sampling variation, we performed permutation tests by randomly subsampling varying proportions of the total species pool at each site. For each level of taxonomic diversity, we resampled species and recalculated the GDIs across 10,000 iterations. We then identified sites where the GDI estimate showed significant deviation from the full-sample expectation. This analysis revealed that only 532 sites (<1%) exhibited statistically significant bias due to variation in sampling proportion, providing additional evidence for the effectiveness of our correction procedures.

#### Simulation Study - Assessing GDI Performance Under Varying Sampling Scenarios

To further validate the behavior of GDI under controlled conditions, we conducted a simulation study to explore the effects of species richness *(S)* and sampling proportion $S/S_{total}$ on GDI accuracy. We generated a synthetic genetic diversity matrix representing 100,000 virtual sites, each with varying numbers of species *(S)* and known "true" nucleotide diversity values per species. For each site, different sampling proportions were randomly drawn, and the corresponding GDIs were calculated using the same correction methods as applied in our empirical data (see Appendix Fig. S3&S4 for output plots). The difference between the true and estimated GDI was quantified using the relative error (RE) calculated as:

$$RE = \frac{abs(cGDI - true\ mean)}{true\ mean}$$

where cGDI is the estimated mean genetic diversity index from the sampled data, and true mean is the actual mean nucleotide diversity of all species at the site.

These errors were then plotted as a function of species richness *(S)* and sampling proportion $S/S_{total}$ to assess how bias and variance in GDI estimates changed with sampling effort (see main Fig. 3). From our simulation, we found that: i) bias decreased with increasing sampling proportion $S/S_{total}$, confirming that more complete sampling yields more accurate GDI estimates, and ii) variance was higher in low-richness sites but was effectively controlled in by our correction methods, especially with the regression-based adjustment. These simulations provide strong support for the statistical robustness and scalability of our GDI framework under variable sampling conditions.

#### GAM analysis of diversity–sampling relationships

To evaluate the relationship between the Genetic Diversity Index (GDI) and sampling parameters, we fitted Generalized Additive Models (GAMs) using the `mgcv` package in R (Wood, 2017). GAMs extend generalized linear models by allowing non-linear relationships between predictors and the response through smooth functions. The model used the form:

$$\text{GDI} = \beta_0 + s(X) + \varepsilon,$$

where $s(X)$ is a smooth function of either species richness ($R$) or sampling intensity ($s$), $\beta_0$ is the intercept, and $\varepsilon$ is the residual error. Thin-plate regression splines were used with the smoothing parameter estimated via restricted maximum likelihood (REML).

Model performance was summarized using three statistics:

1. **Deviance explained**:
$$D_{\text{expl}} = 1 - \frac{D_{\text{model}}}{D_{\text{null}}},$$
where $D_{\text{model}}$ and $D_{\text{null}}$ are the model and null (intercept-only) deviances, respectively. This value represents the proportion of total variation accounted for by the GAM, analogous to the coefficient of determination ($R^2$) in linear regression. Values close to 1 (or 100 %) indicate that the model explains most of the variation in GDI, while values near 0 indicate weak explanatory power.

2. **GAM $R^2$** ($R^2_{\text{GAM}}$): This statistic approximates the proportion of variance in the response explained by the fitted smooths for Gaussian models, providing a direct analogue to the adjusted $R^2$ of a linear regression.

3. **Effective degrees of freedom (edf)**: The edf quantifies the complexity of each smooth term. An edf of 1 indicates an approximately linear relationship, whereas values greater than 1 denote increasing

non-linearity or curvature in the fitted trend. Similar edf values (e.g., around 3) across metrics indicate comparable degrees of smoothness, suggesting consistent levels of non-linearity.

Ninety-five percent confidence ribbons around the fitted GAM curves represent the uncertainty of the predicted mean response, computed as twice the standard error of the fitted values.

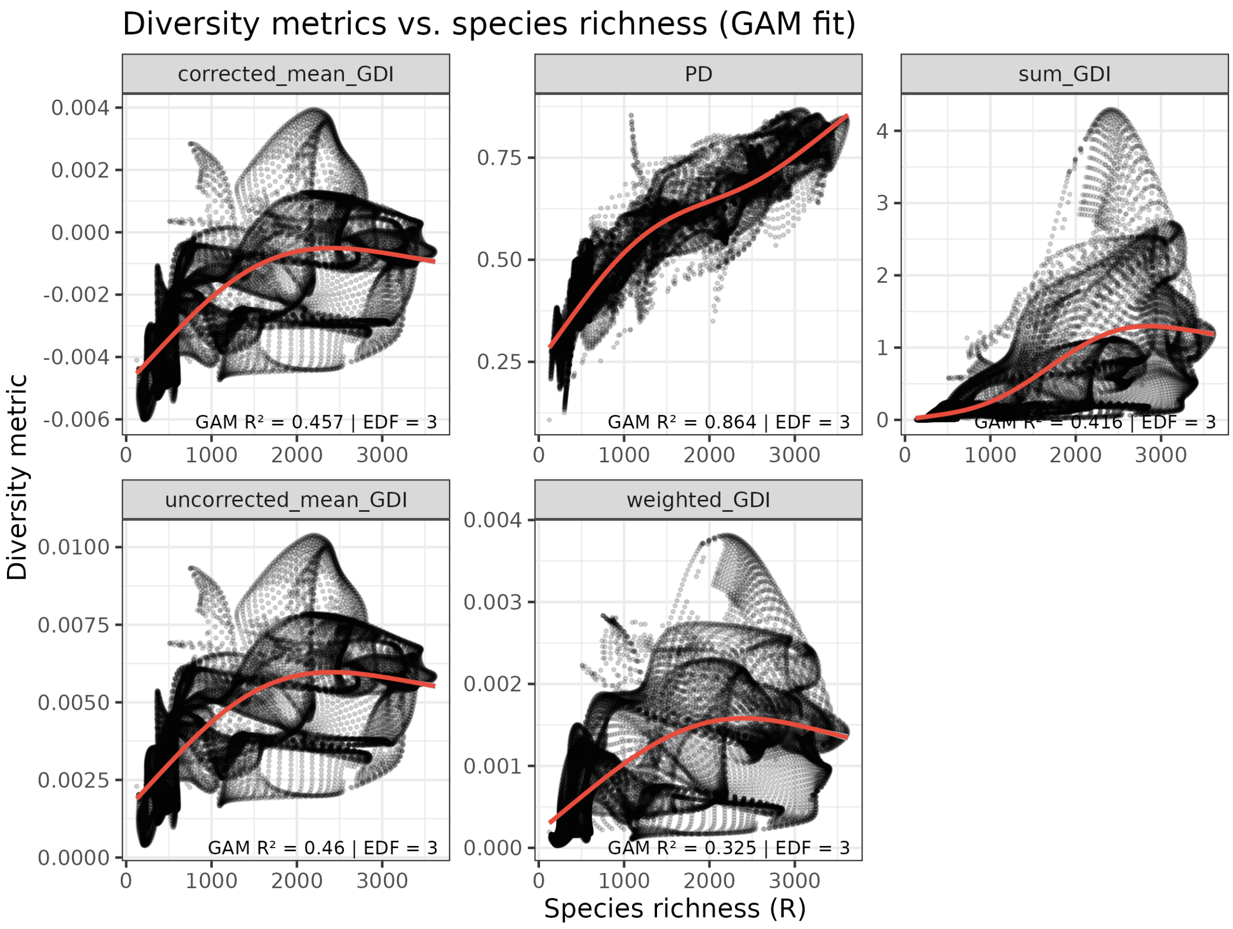


Figure S3: GAM analysis showing the relationship between uncorrected GDI and sampling proportion (number of species sampled/ total species richness) across Europe. A. Mean Genetic Diversity Index (mGDI), B. Sum Genetic Diversity Index (sGDI). The plots show the fitted GAM curves (black line) with 95% confidence intervals (grey ribbon). The deviance explained, GAM R2, and effective degrees of freedom (edf) are indicated in each panel. Linear relationships between four versions of the Genetic Diversity Index (GDI) and phylogenetic diversity (PD) with species richness, calculated across 1,860 vascular plant species in Europe. Each panel shows a linear regression fit between species richness and a diversity metric. The adjusted R2 values indicate the proportion of variance in each diversity metric explained by species richness, accounting for the number of observations and model complexity. This provides a measure of how strongly species richness predicts each type of diversity.

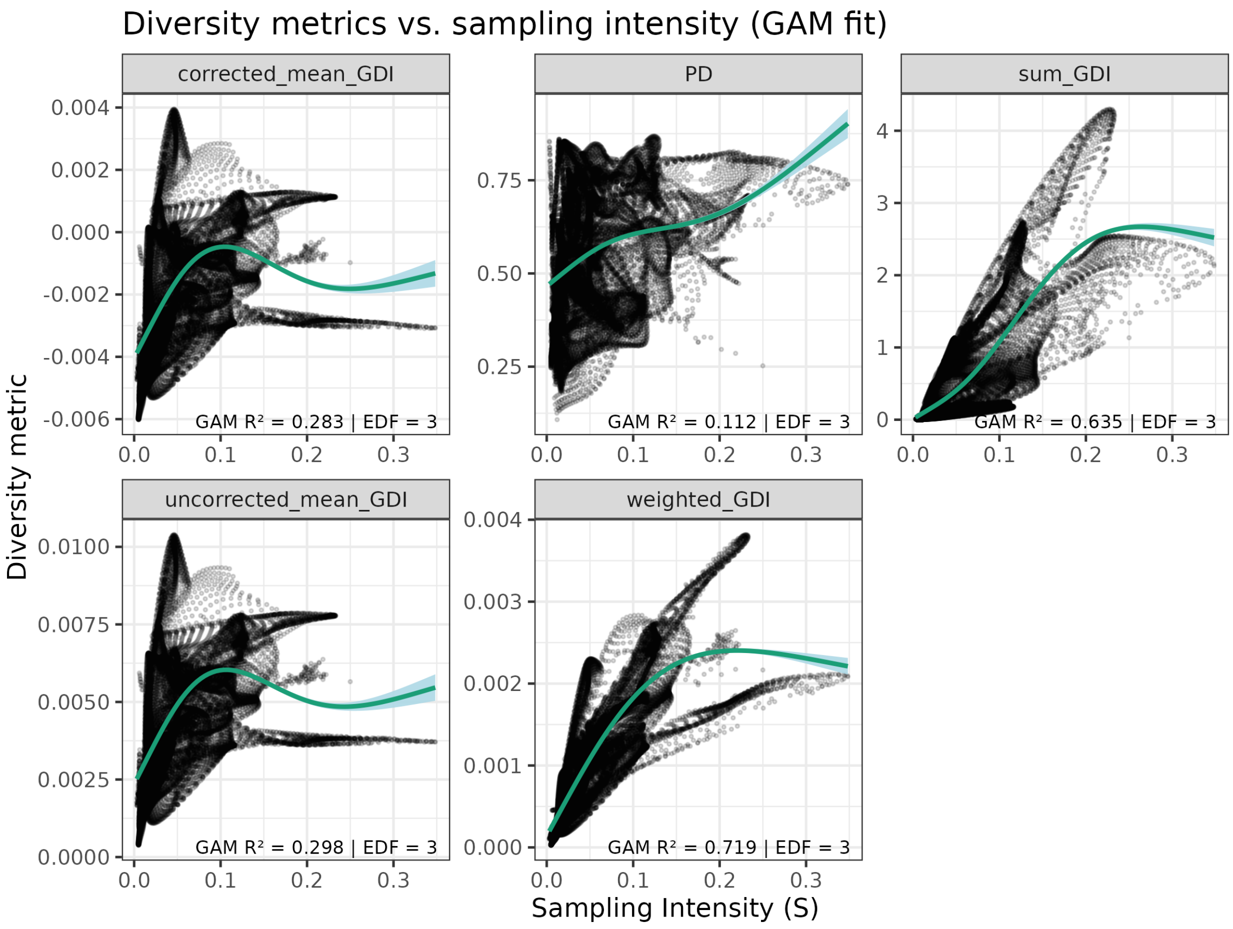


Figure S4: Linear relationships between four versions of the Genetic Diversity Index (GDI) with sampling proportion, defined as the proportion of species sampled relative to the total number of species occurring in each raster cell, calculated across 1,860 vascular plant species in Europe. Each panel presents a linear regression between sampling proportion and a diversity metric. The adjusted R2 values indicate the proportion of variance in each diversity metric explained by sampling proportion, while accounting for the number of observations and model complexity. This illustrates how sensitive each diversity metric is to variation in sampling effort.

Table S1: Linear model explaining standardized Genetic Diversity Index (GDI) as a function of climate stability and sampling intensity. Predictors are z-transformed.

| Predictor | Estimate | SE | t | *p* |
|---|---|---|---|---|
| Intercept | 0.000 | 0.004 | 0.00 | 1.00 |
| Temperature stability | 0.480 | 0.005 | 106.53 | $< 2 \times 10^{-16}$ |
| Precipitation stability | 0.306 | 0.005 | 66.15 | $< 2 \times 10^{-16}$ |
| log(Sampling intensity) | -0.213 | 0.005 | -44.47 | $< 2 \times 10^{-16}$ |

#### Mapping Genetic Diversity

All spatial processing and mapping were performed in R using a WGS 84 (EPSG:4326) coordinate reference system. The final output raster has a resolution of 0.25° × 0.25° (approximately 27.8 km pixel edge length at the equator, narrowing to ~14–25 km across the European map range of 20°N–85°N) and covers a spatial extent of -10°E to 70°E longitude and 20°N to 85°N latitude (260 rows × 320 columns). Rasterization and

Table S2: Generalized additive model (GAM) explaining standardized GDI. Spatial autocorrelation is accounted for using a two–dimensional smooth of coordinates.

| Parametric term | Estimate | SE | t | $p$ |
|---|---|---|---|---|
| Intercept | 0.000 | 0.001 | 0.00 | 1.00 |
| Temperature stability | -0.001 | 0.005 | -0.12 | 0.91 |
| Precipitation stability | 0.032 | 0.005 | 6.91 | $4.9 \times 10^{-12}$ |
| log(Sampling intensity) | 0.165 | 0.012 | 13.68 | $< 2 \times 10^{-16}$ |

| Smooth term | edf | F | $p$ |
|---|---|---|---|
| s(x,y) | 298.4 | 3151 | $< 2 \times 10^{-16}$ |

spatial analysis were carried out using the packages terra (Hijmans, 2025), sf (Pebesma, 2018), and gSoup (Karunarathne, 2024), while community-level phylogenetic diversity computations were managed with the phyloregion package (Daru et al., 2020). Resulting maps were visualized using ggplot2 (Wickham, 2016).

### A Global Outlook for Genetic Diversity Assessment

#### Data Availability and Feasibility

We assessed the global applicability of the Genetic Diversity Index (GDI) using a dataset of one million geo-referenced accessions across major vascular plant groups. This revealed broad taxonomic representation in well-sampled regions like Europe, North America, Southeast Asia, and Australia, with both ecologically diverse and economically important taxa included (see Fig. S5).

However, significant data gaps persist in South America, Africa, Western Asia, and Siberia. These disparities reflect uneven research investment and infrastructure, where biodiversity-rich regions often lack sufficient sequencing coverage. Despite this, our simulations show that GDI remains robust when a taxon-specific threshold of data sufficiency is met (see main text Fig.3). This implies that complete datasets are not required and targeted sampling and adaptive thresholds can support accurate GDI assessments even with partial coverage.

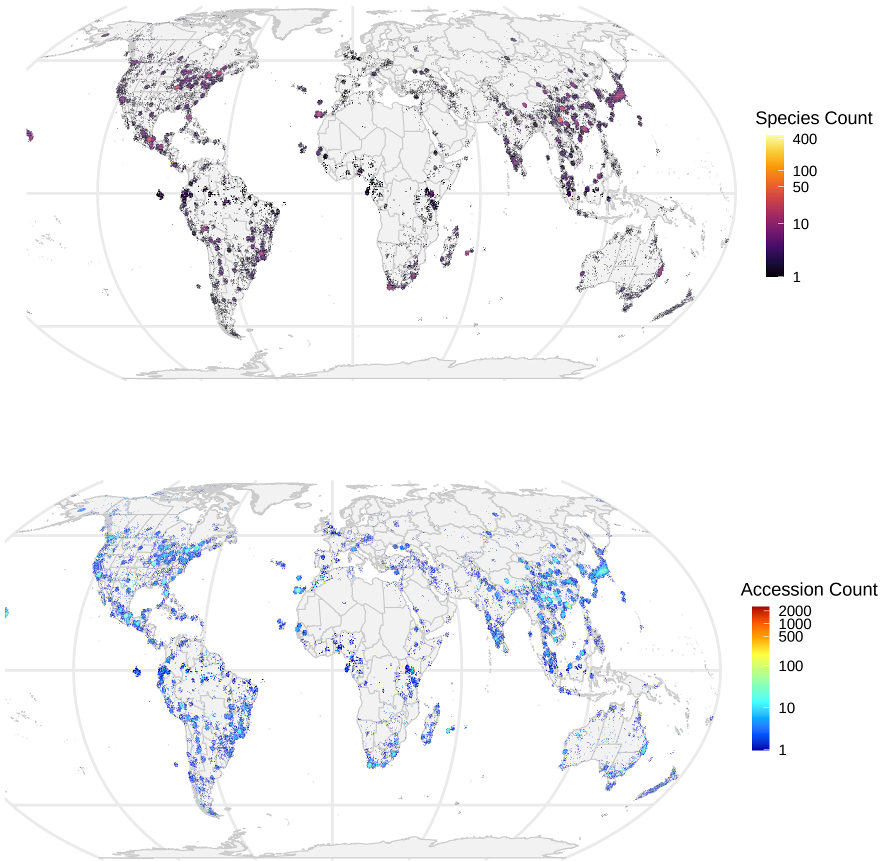


Figure S5: A global map showing the distribution of georefenced genetic data across major vascular plant groups processed from **one million accessions** from NCBI GeneBank. The top panel shows the number of species with genetic data, while the bottom panel shows the number of accessions per species. The map highlights the uneven distribution of genetic data availability, with regions such as Europe and North America having higher representation compared to South America and Africa. This underscores the need for targeted efforts to enhance genetic data coverage in underrepresented regions, particularly in biodiversity hotspots. Color gradients are on a log-scale to improve the visibility.

However, challenges remain in standardizing metadata quality, ensuring taxonomic accuracy, and addressing sampling biases. Future efforts should focus on improving data curation, expanding sequencing initiatives in underrepresented regions, and integrating genetic data with ecological and environmental datasets to enhance the utility of GDI for global biodiversity assessments. We discuss these considerations in detail in the main text.

### Appendix 2: META-DATA ANALYSIS OF GENBANK SEQUENCES FOR EUROPEAN VASCULAR PLANTS

This section summarizes the results of our meta-data analysis of publicly available genetic sequence data for European vascular plants, sourced from the NCBI GenBank. The full methodology is detailed in the Appendix 1 Methods section.

#### NCBI Nucleotide Database Exploration

Our analysis revealed a total of 24,569,450 sequence accessions associated with vascular plant species native to Europe. Of these, 280,979 accessions (1.2%) contained primary geo-referenced information, i.e., geographic coordinates embedded in the metadata fields. In addition, 2,772,037 accessions (11.7%) included secondary geo-referenced information, consisting of place names or location descriptors. We applied geocoding procedures to the secondary metadata and successfully retrieved coordinates for 2,737,670 accessions (98.8%), whereas 34,367 accessions (1.2% of secondary records) could not be resolved to usable geographic coordinates. Taken together, approximately 13% of the total accessions included either direct or derived geo-referenced information and were considered suitable for spatial analysis (Appendix Fig. S6A).

#### Data Coverage Across Taxonomic Levels

Among the 26,242 native vascular plant species recognized for Europe, 9,409 species (35.9%) were represented in GenBank by at least one sequence. Within this sequenced subset, 2,827 species (29%) were linked to accessions with primary geo-referenced metadata. These species spanned 134 of the 154 native plant families (87%) and 812 of the 1,267 genera (64%). In contrast, 7,143 species (72%) had accessions containing secondary location metadata, which, after successful geocoding, extended coverage to 148 families (96%) and 1,161 genera (91%). When combining species with either primary or geocoded secondary coordinates, a total of 7,512 species (approximately 77% of all sequenced species) could be associated with geographic coordinates. This combined set included 1,189 genera, representing 92% of the native European flora at the genus level. Of the geo-referenced species, 6,949 (92.5%) were further linked to usable DNA marker information and were retained for genetic diversity analyses (Appendix Fig. S6B).

#### Taxonomic Biases in Geo-referenced Data

The distribution of geo-referenced data across taxonomic groups revealed pronounced biases. Several large families with high species richness were underrepresented in terms of spatial genetic data. For example, only 12% of the 8,659 native species in Asteraceae were associated with geo-referenced sequence accessions. Similarly low representation was observed in Rosaceae (14% of 1,979 species) and Ranunculaceae (12% of 1,432 species). In contrast, families with fewer species, ranging from 7 to approximately 1,000 native taxa, were generally better represented, with an average of 48% (±2%) of their species linked to geo-referenced sequence data. These patterns suggest a historical sequencing emphasis on taxonomically narrower groups, model species, or economically relevant taxa, leading to significant gaps in coverage for some ecologically prominent families (Appendix Fig. S7).

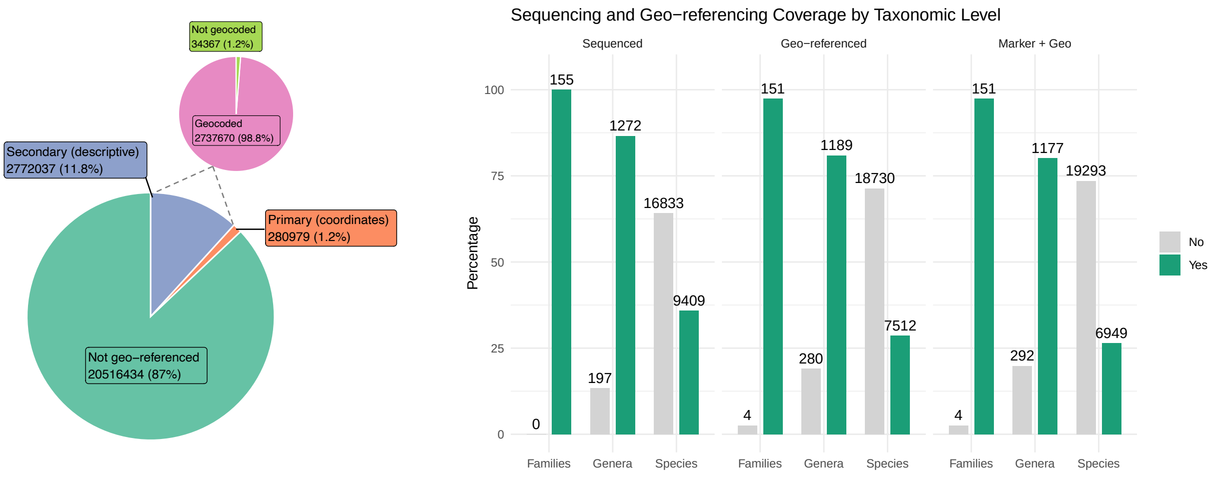


(A) (B)

Figure S6: Data availability of European vascular plants in NCBI GenBank. A. Hit index indicated as the number of accessions, and B. data availability shown at the species, genus, and family levels, representing different taxonomic scales. Hit indices are illustrated using percentage bars and accompanying data tables for three categories: (1) sequenced records, (2) geo-referenced records, and (3) geo-referenced records with usable sequence data (i.e., with markers).

#### Species Representation and Data Coverage

We further assessed the geographical distribution of species richness across Europe using data from the World Checklist of Vascular Plants (WCVP). Species richness varied markedly across regions, with as few as 188 species in Svalbard and up to 9,865 species in Türkiye. Spain exhibited the highest species richness in Western Europe, with 6,312 native species, and overall richness was concentrated in the Mediterranean floristic regions (Appendix Fig. 3). These patterns differ from those observed in species distribution maps derived from occurrence-based datasets (e.g., Daru, 2024, SDM), which are restricted to species with verified spatial records.

#### Spatial Distribution of Geo-referenced Genetic Data

To evaluate sequencing depth, we analyzed the number of accessions per species. Species-level coverage across European regions varied depending on the minimum threshold of sequences per species. With a threshold of at least one sequence, regional coverage ranged from 37.3% to 88.6% (mean: 71.3%). Increasing the threshold to ten sequences per species reduced mean coverage to 56.1%, while applying a threshold of 100 sequences per species further reduced coverage to 16.6% (Appendix Fig. S9). A similar trend was observed in geo-referenced sequence data. Taxonomic coverage was 43.6% for primary geo-referenced data and 61.1% for secondary data, increasing slightly to 62.1% when the two were combined. However, coverage declined consistently with higher sequence thresholds, illustrating a trade-off between sequence depth and taxonomic breadth (Appendix Fig. S8).

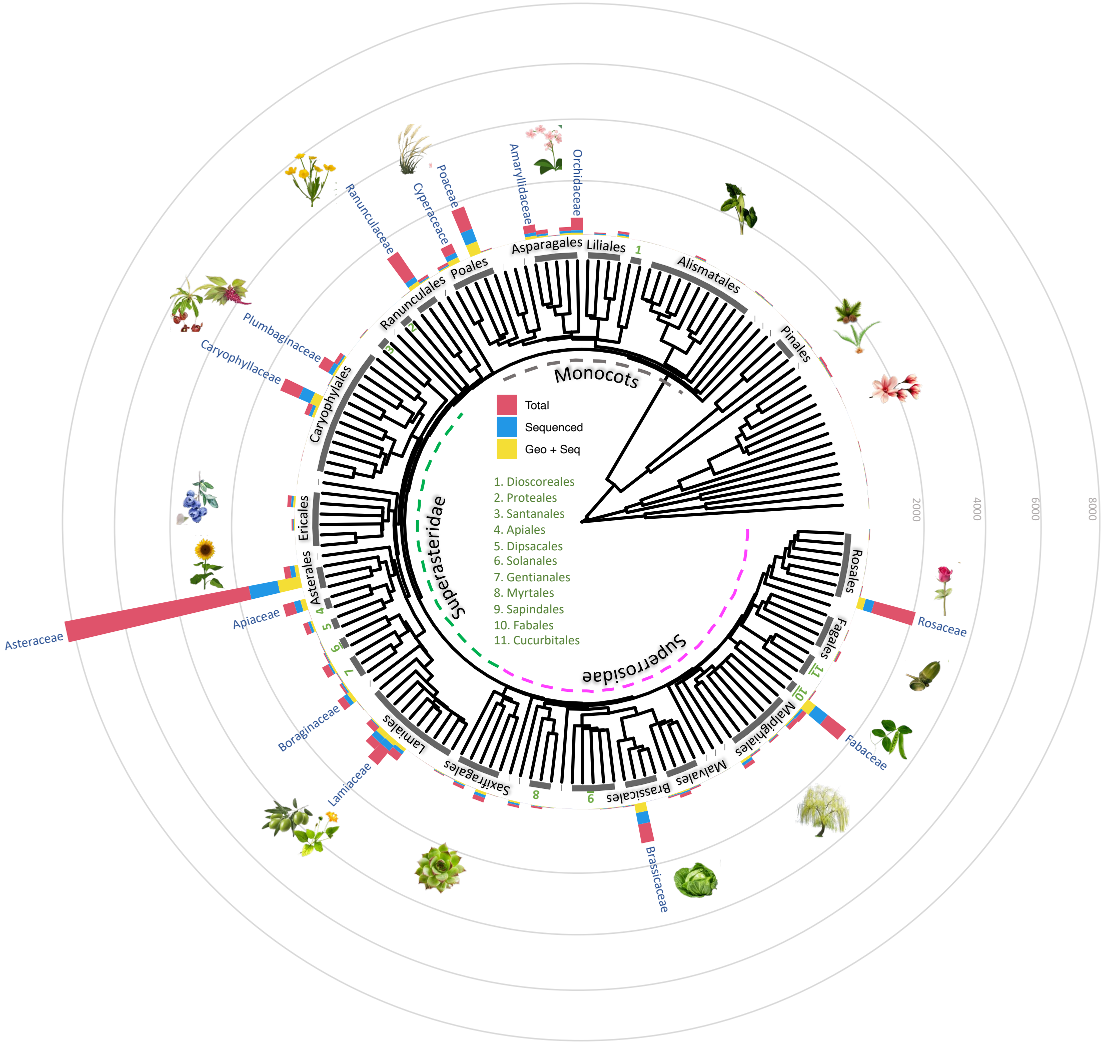


Figure S7: Phylogenetic tree of European vascular plant families, showing the representation of sequenced and georeferenced accessions in NCBI. The red bar shows the total number of species in a family occurring in Europe, blue bar shows how many species of them have been sequenced, and the yellow bar shows how many of those sequenced species have georeference. Major vascular plant families in Europe have been labeled in blue font. Plant orders are labeled between the tree and the barplots. The family level phylogenetic was built using the R package V.Phylomaker2 (Jin and Qian, 2019), which uses an extended version of GBOTB tree from Smith and Brown (2018).

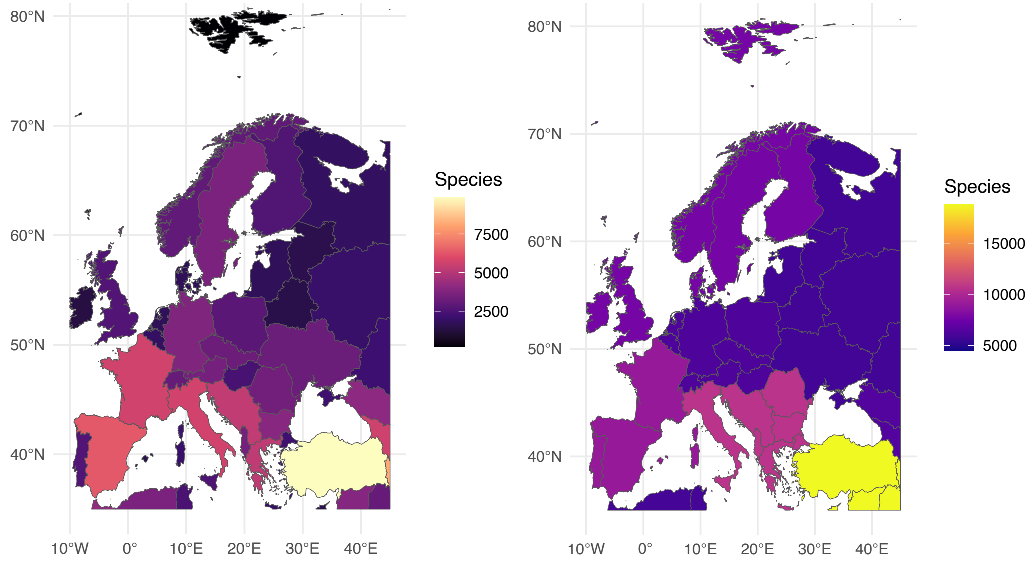


(A) (B)

Figure S8: Complete species richness (number of species) in the European Region according to World Checklist of Vascular Plants consisting of 26,242 species, summarized by TDWG (Brummitt et al., 2001) A. level 3 (Floristic country) and B. level 2 (Floristic Region).

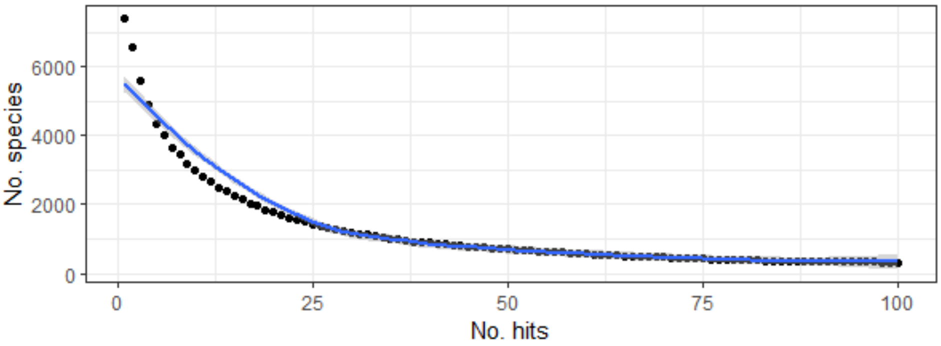


Figure S9: NCBI nucleotide database coverage of vascular plant Flora in Europe for combined geo-referenced sequences with marker information. Line plot shows the number of species versus number of accessions (hits) with geo-referenced sequences (black: number of species blue LOESS smooth line to show the trend).

#### Combined Geo-referenced Data with Markers

After filtering for both marker information and spatial data, 629,175 accessions ( 2.7% of 23,375,684 unique accessions) remained. These accessions were associated with 6,947 species and covered a wide geographic range. The distribution of these sequences across a  450 km2 raster cell grid revealed strong spatial heterogeneity, with an average of 134 sequences per grid cell. Grid cell values ranged from one to 2,138 sequences, with hotspots centered around Antwerp (Belgium), Prague (Czech Republic), Ankara (Türkiye), and Madrid (Spain) (Appendix Fig. S10).

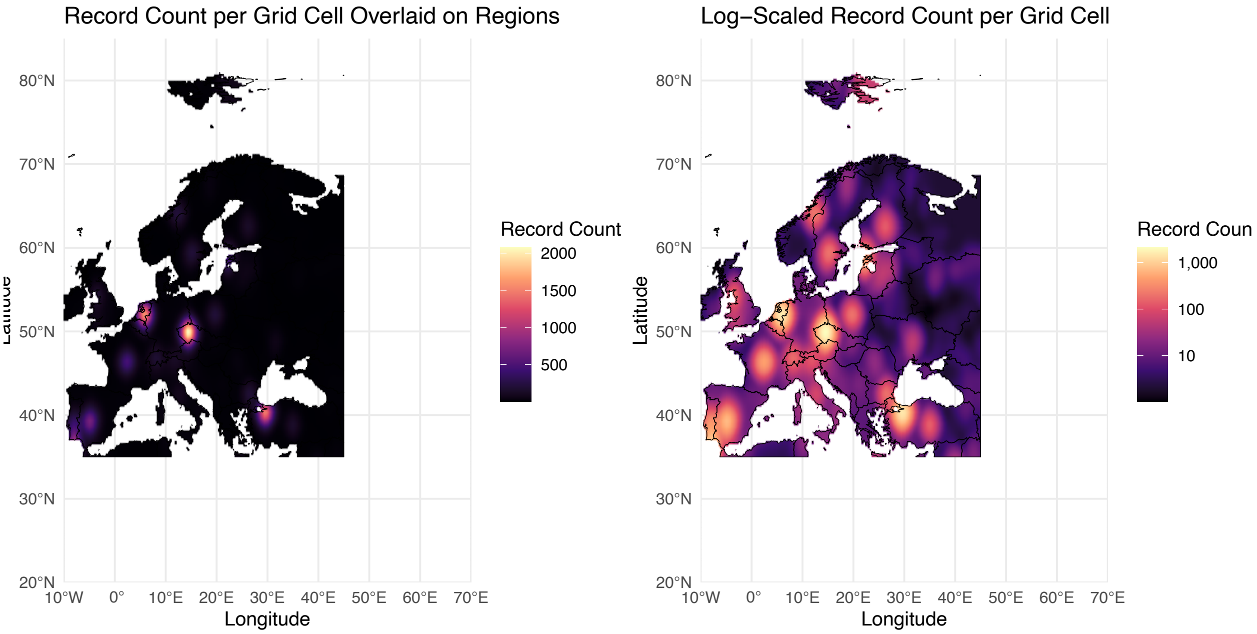


Figure S10: Geographical distribution combined geo-referenced data. To visualize the geographical distribution of data, all sequences were plotted on a 400 km2/cell grid and counted. A. Combined geo-referenced data with markers on a grid. B. Combined geo-referenced data with markers in a raster grid on a logarithmic scale. Each grid-cell indicates the number of sequences from high (yellow) to low (purple).

#### Combined Geo-referenced Data on a Threshold

To refine the dataset, thresholds were set to ensure broad geographical coverage and accurate nucleotide diversity calculations. Filtering the combined geo-referenced data with marker information for thresholds of at least five hits per species-marker combination resulted in 623,721 sequences. These sequences represented 2,468 species (33%) out of the 7,512 species available for geo-reference and marker information. Therefore, downstream analysis focused on a dataset containing species with at least five hits per species-marker combination. Appendix Figure S11 shows the number of species and accessions used in the analysis summarized for TDWG floristic countries of Europe.

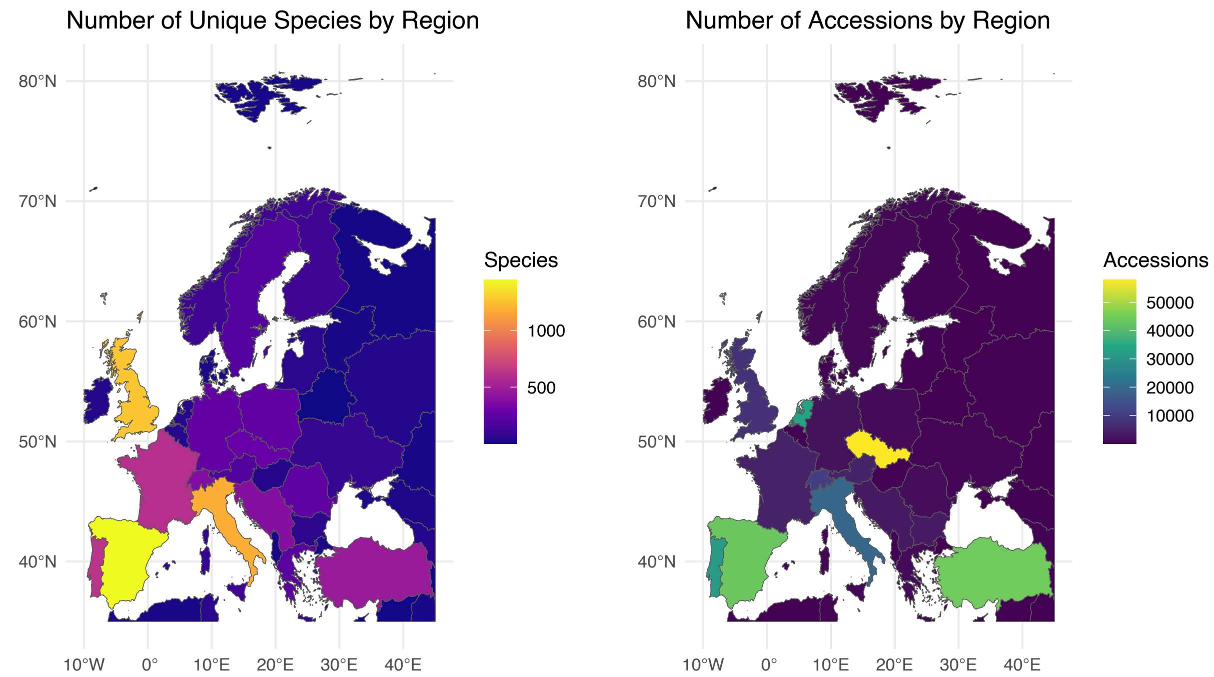


Figure S11: Distribution of georeferenced DNA sequences used in the genetic diversity analysis in this study summarized by A. unique species and B. number of accessions, for TDWG level 3 (floristic country).
